# Resolving the treachery of images: Neural signatures during natural interactions with real objects are as reliable as while viewing images

**DOI:** 10.64898/2026.04.12.717765

**Authors:** Sini Simon, Surbhi Munda, Thomas Cherian, Shubhankar Saha, Georgin Jacob, Jhilik Das, Debdyuti Bhadra, SP Arun

**Affiliations:** Centre for Neuroscience, Indian Institute of Science, Bangalore 560012

## Abstract

Is a real object the same as its image? Vision and its neural basis have been studied almost entirely with images. However, real objects are fundamentally different, because *we can act on them*. But since we see and interact with real objects differently each time, it is widely believed that brain activity will be highly variable. An alternative view is that real objects should evoke reliable neural signatures since brain activity might be optimized for natural behavior. To investigate these possibilities, we performed wireless neural recordings from the high-level visual (IT) and motor regions (PMv/PFC) of freely moving monkeys, performing a natural task where they ate food items offered by humans, and a controlled task where they viewed the corresponding images on a screen. Neural decoders trained on images accurately identified the corresponding real objects. Even though neural responses were more variable during real object interactions, the variability of object-relevant information was strikingly similar, suggesting that the additional variability is present along irrelevant dimensions. Finally, key events in the natural task elicited reliable neural signatures. We conclude that brain activity is efficiently organized during natural behavior, ensuring that real objects are encoded as efficiently as images.

## INTRODUCTION

Real objects are qualitatively different from their images, as famously noted by Rene Magritte in *The Treachery of Images (1929)*. Indeed, real objects contain far richer cues like stereo, motion and parallax – and more importantly, *we can act on them* (Snow and Culham, 2021; Culham and Deligiannis, 2026). However, nearly all our understanding of vision and its neural basis comes from studying images, because images are easy to control and obtain repeatable responses (Culham and Deligiannis, 2026). Studying real objects on the other hand is complicated, because each interaction with them can be different: even the simple act of eating can involve seeing and manipulating the food differently each time (Müller and Sternad, 2009; Dhawale et al., 2017).

Do real objects and their images evoke similar or different neural activity? This question might seem trivial at first, but there are two deeper issues that merit careful consideration. First, there is wide variation in how animals and even humans react to real objects compared to images (Bovet and Vauclair, 2000). This difference could arise from qualitatively different underlying neural representations in the visual cortex, or alternatively in motor regions that generate different affordances to real objects and images (Culham and Deligiannis, 2026). At the neural level, some (but not all) neurons in high-level visual areas respond similarly while viewing real objects and their images (Tanaka et al., 1991; Kobatake and Tanaka, 1994), but many neurons are also sensitive to stereo disparity signals present only in real objects and not in images (Connor et al., 2007; Orban, 2011). Thus, there is reason to expect similarities and differences. However, these results are based on neural recordings performed while animals are constrained from making movements or interacting with these objects. As a result, we know very little about how neurons in high-level visual regions respond during natural behaviors while animals interact freely with real objects.

Second, are neural responses more reliable or less reliable while interacting with real objects compared to viewing images? There is an ongoing debate regarding the nature of neural activity during natural behavior and its relation to more controlled settings. The prevailing view is that neural activity will be less reliable since natural behaviors involve large sensory and motor variations (Miller et al., 2022; Cisek and Green, 2024; Leopold, 2024). Vision is thought to be even more variable since natural viewing also involves eye movements that introduce neural variability (Xiao et al., 2024; Singh et al., 2025). Alternatively, brain activity might be optimized for natural behaviors, in which case the brain must generate neural representations that are systematic and reliable despite the variability inherent in natural behavior (Miller et al., 2022; Parodi et al., 2025). We reasoned that such reliable representations are likely to be in higher order visual and motor regions that are known to contain invariant representations (Rizzolatti and Luppino, 2001; Connor et al., 2007; DiCarlo et al., 2012; Conway, 2018; Schrijver et al., 2024). Addressing this issue will require recording brain activity under both controlled and natural settings as proposed recently (Cisek and Green, 2024; Parodi et al., 2025).

### Overview of this study

Here, we set out to investigate these fundamental gaps in our knowledge through wireless brain recordings in freely moving monkeys in controlled and natural conditions. We recorded simultaneously from the inferior temporal (IT) cortex, which contains invariant visual object representations (Connor et al., 2007; DiCarlo et al., 2012; Conway, 2018) and from the premotor/prefrontal regions (ventral premotor cortex, PMv and ventrolateral prefrontal cortex, PFC), which contain invariant motor and task representations (Rizzolatti and Luppino, 2001; Schrijver et al., 2024). In the natural setting, we recorded brain activity while monkeys housed in a large arena took food items offered by human experimenters (Figure 1A). In the controlled setting, we recorded brain activity while monkeys performed a fixation task involving images of the same food and experimenters on a screen (Figure 1B). This allowed us to carefully compare and interpret the complex neural signatures in the natural task by relating them to neural responses to images in the more controlled setting.

**Figure 1.**
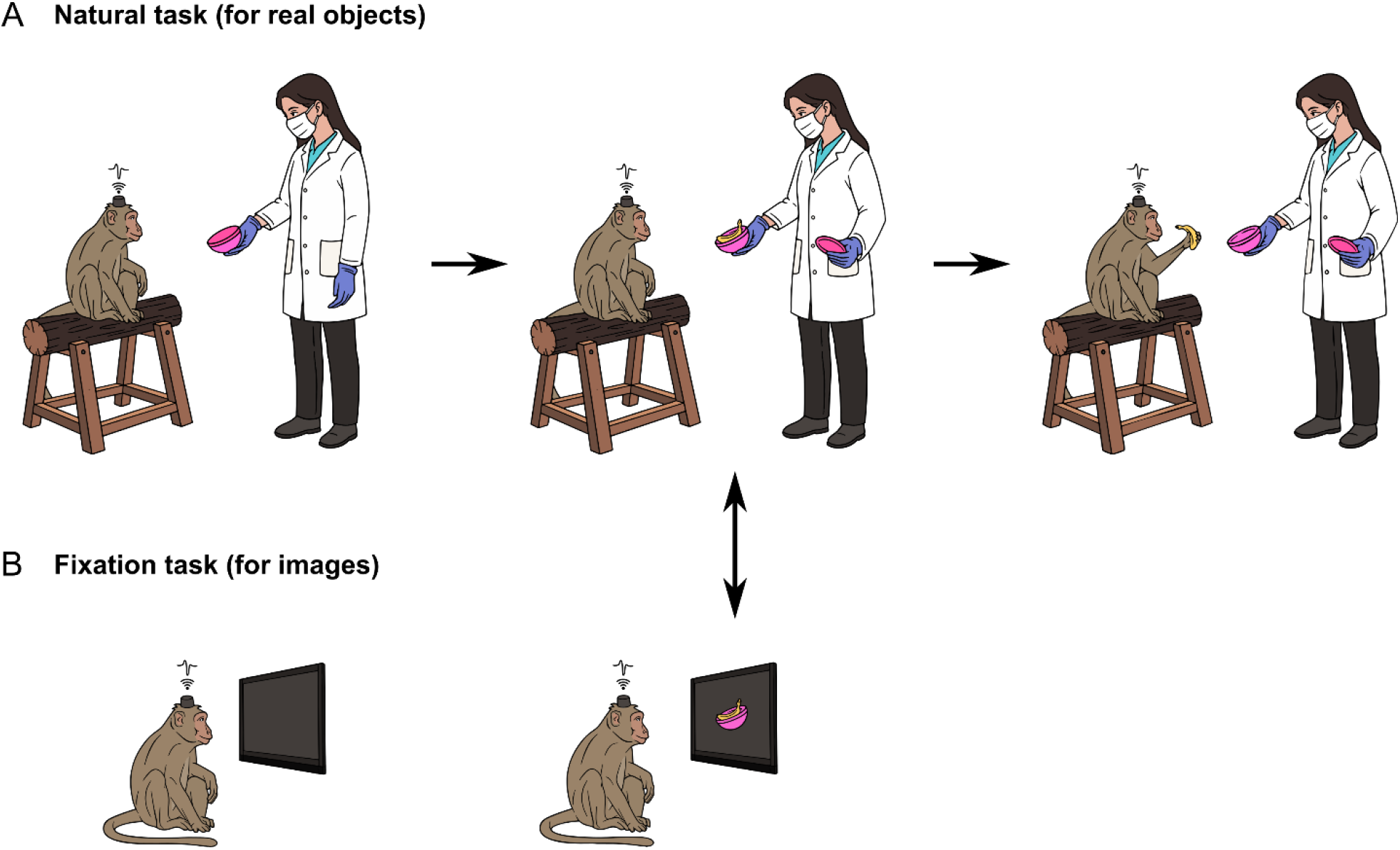
Comparing neural activity to real objects and their images. (A) In the natural task, a human experimenter presented a box with a closed lid to the monkey (*left*), then opened the lid to reveal food (*middle*), after which the monkey took and ate the food (*last*). Across trials, there were two human experimenters (E1/E2) and three types of food (banana/carrot/raisin). The natural task was designed so that the opening of the lid created a visual event that was as closely matched as possible to the onset of images in the fixation task, to facilitate comparison of neural activity between the two tasks (*indicated by arrow*). (B) In the fixation task, the monkey viewed images of food and faces in multiple views, presented at a size similar to what they would encounter in the natural task.

## RESULTS

Each monkey participated in two sessions. In the natural session, one of two human experimenters entered the room, approached the monkey holding a box with a closed lid, opened the box in front of the monkey to reveal one of three types of food (banana, carrot or raisin pieces), which the monkey took and ate, after which the experimenter left the room (Figure 1A). We designed this task so that there was a specific time at which the food was revealed to the monkey, which can then be compared with neural responses when the monkey viewed the same images (Figure 1B). In the touchscreen session (Figure 1B), the monkey had to perform a fixation task involving a series of images to characterize visual responses. The monkey also performed a motor task with its left and right hand separately to characterize motor responses.

For the natural task, each monkey was tested in a large natural arena enriched with tree branches, perches, ropes and swings (Figure 2A). This arena was connected through a series of movable partitions to a room containing a touchscreen workstation with gaze tracking abilities (Figure 2B), as described previously (Jacob et al., 2021). We performed wireless brain recordings from 256 electrodes implanted into the higher order visual and motor regions (IT, PMv, PFC) of two monkeys (Figure 2C; see Methods).

**Figure 2.**
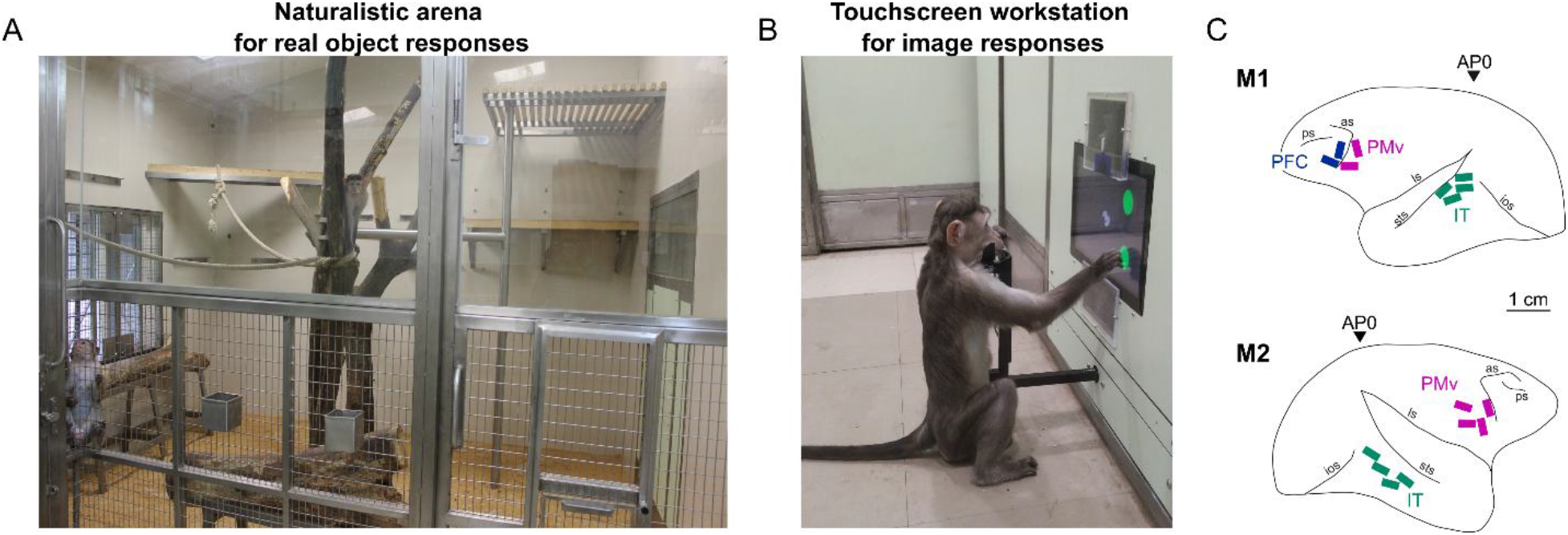
Wireless brain recordings in natural and controlled settings. (A) Brain activity during natural behavior was recorded wirelessly in a large natural arena (3.3 x 4.13 m) enriched with perches, tree trunks, and ropes in which monkeys were group-housed throughout the study and separated after electrode implantation. For the natural task, each monkey was separated from its partners using a series of partition doors, one of which is partially visible at the extreme left. (B) Brain activity in controlled settings was obtained using a touchscreen workstation with gaze tracking capability, where each monkey performed a fixation task. The image shows a naïve monkey performing a task. (C) Electrode array locations in each monkey, reconstructed by aligning sulcal landmarks in MRI scans with photographs of the surgically implanted arrays. Monkey M1 had 128 electrodes in IT, 64 in PMv and 64 in PFC. Monkey M2 had 128 electrodes in IT & 128 in PMv. The inverted triangles indicate the stereotaxic zero (AP0).

### Neural decoding of visual and motor information in the touchscreen task

Since brain activity during natural behavior might be complex and variable, we first set out to characterize the visual and motor information present in the controlled settings, while animals performed a visual task and a motor task (see Methods).

In the visual task, animals had to fixate at series of images of food, faces and arena (Figure 3A). Figure 3B depicts example neural responses recorded in monkey M2 from all 128 electrodes of IT in the fixation task. It can be seen that many channels showed phasic activity that was time-locked to the visual stimulus. In the motor task, the monkey had to touch a hold button, and then make movements to one of four locations on the screen with his left or right hand in separate blocks (Figure 3C). Figure 3D depicts example neural responses from all 128 electrodes of PMv in the left and right hand motor tasks. It can be seen that many channels showed phasic activity that was time-locked to the movement onset in the motor task (for PMv). In this manner, we identified a total of 190 visual neurons in IT (92 of 128 from M1, 98 of 128 from M2), 93 movement neurons in PMv (37 of 64 from M1, 56 of 128 from M2) and 16 visual/movement neurons in PFC (16 of 64 from M1) for further analyses.

**Figure 3.**
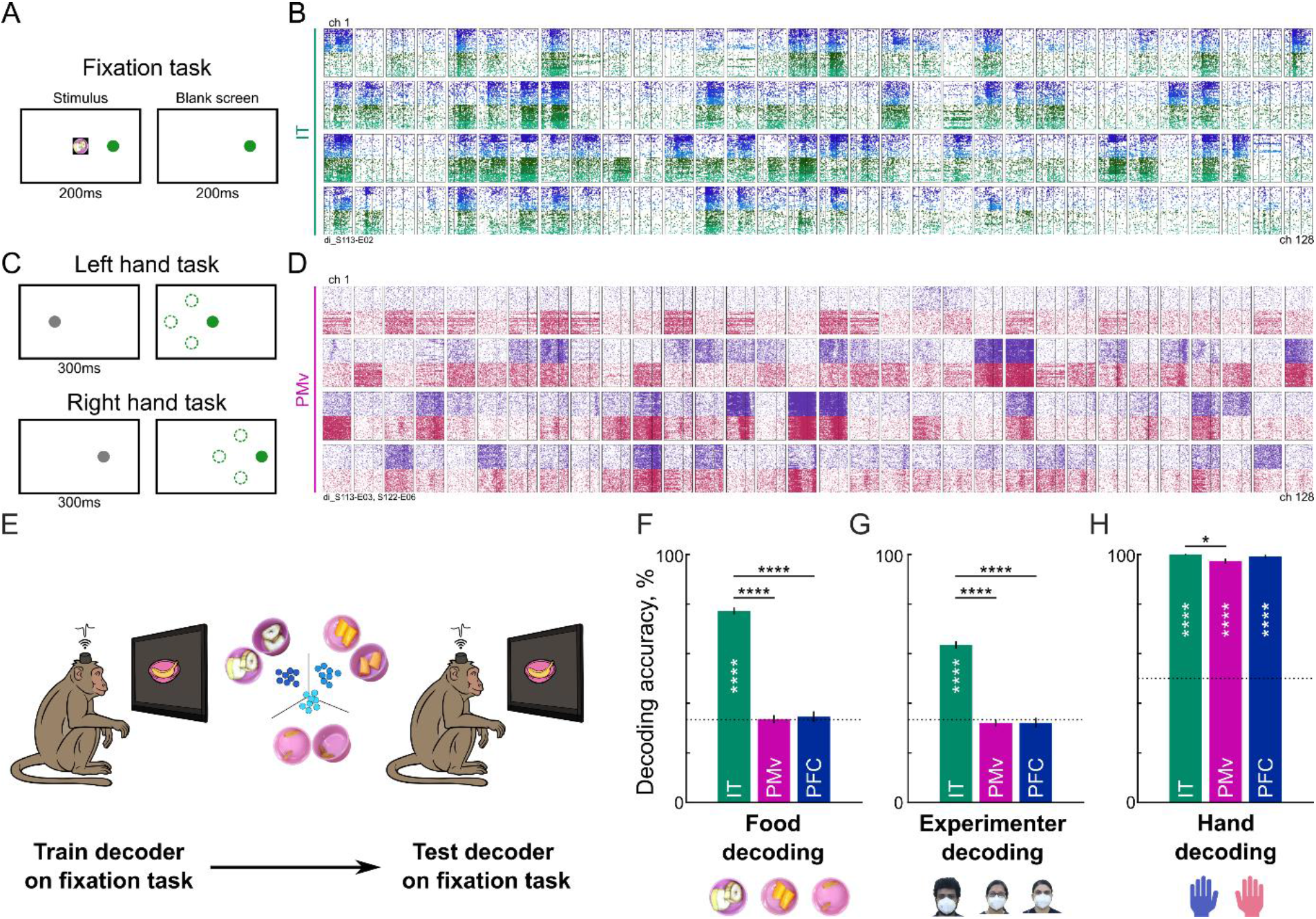
Neural decoding of visual and motor information in touchscreen tasks. (A) Schematic of the fixation task. On each trial, the monkey had to touch the green button, and then had to fixate on a series of images that appeared on the screen for 200 ms with a 200 ms blank interval, and received a juice reward for maintaining fixation throughout. Images are shown against a white background here, but were shown against a black background in the actual experiment. (B) Example multiunit activity recorded from all 128 electrodes in IT cortex of monkey M2. Each row shows the multiunit activity from 32 electrodes, with individual boxes containing rasters from one electrode. Black lines within each box represent the image on and off times. Individual rows of ticks within each box represent the spikes detected from single trials of food images (*blue shades: dark is banana, light is carrot, lightest is raisin; ∼16 correct trials/food*) or experimenter images (*green shades: dark is E1, light is E2, lightest is E3; ∼16 correct trials per image*). A total of 190 of 256 electrodes (M1 & M2 combined) showed clear visual responses to these images in IT. (C) Schematic of the left and right hand tasks. On each trial, the monkey had to hold a button to start the trial, and then touch one of four target buttons that appeared after a brief delay. Images in the actual experiment are shown against a black background. (D) Example multiunit activity recorded from all 128 electrodes of PMv cortex of monkey M2, following the same layout as in panel (B). Individual rows of ticks within each box represent the spikes aligned relative to response onset (*black line*) from individual trials (*purple*: left hand, *red*: right hand trials). A total of 93 of 192 electrodes in PMv (M1 & M2 combined) and 16 of 64 electrodes in PFC (in M1) showed clear motor responses aligned to movement onset. (E) Schematic of decoding from touchscreen tasks. We trained linear decoders on the population activity obtained from individual trials of the touchscreen tasks evoked to each food type across views, and tested the decoder on unseen trials of these items. We followed a similar procedure for experimenter and hand decoding. (F) Cross-validated decoding accuracy for food type from IT (*green*), PMv (*purple*) and PFC (*blue*), depicting the mean ± s.e.m across trials. Decoding accuracy was obtained through five-fold cross-validation from separate decoders trained on the population activity in each monkey, and pooled across trials in both monkeys. Asterisks within each bar indicate the statistical significance of decoding from that region (**** is p < 0.00005, chi-squared test comparing observed correct/incorrect counts to that predicted by chance). Asterisks across pairs of bars represent the statistical significance of the comparison between brain regions (**** is p < 0.00005, *** is p < 0.0005, rank-sum test comparing decoder output across trials). (G) Same as (F) but for decoding of experimenter identity. *[Face images are those of the authors D.B., S.M. and S.S]* (H) Same as (F) but for hand decoding in the motor task.

#### Neural decoding of food & experimenter identity in the fixation task

In the fixation task, monkeys viewed three types of familiar food (raisins, banana and carrot pieces) shown in a pink box at multiple views, which they would subsequently interact with in the natural session. They also saw images of three experimenters (with and without face masks) at multiple views, two of whom would subsequently offer food to them in the natural session. We therefore investigated whether neural activity patterns evoked by these images could be used to decode the identity of the food and experimenter in a view-invariant manner, as might be required in a more natural setting. To this end, we trained a linear population activity decoder on neural responses evoked in each brain region (IT/PMv/PFC) during single trials in the fixation task to decode the food item (banana/carrot/raisin) or the experimenter (E1/E2/E3), and tested the decoder on unseen held-out trials (Figure 3E; see Methods). For brevity, this decoder accuracy was pooled across trials in both monkeys, but we obtained similar results in each monkey separately. We found that IT neurons (but not PMv or PFC) reliably encoded both food type (Figure 3F) as well as experimenter identity (Figure 3G). Thus, food type and experimenter identity can be decoded reliably mainly from IT during the fixation task, consistent with previous studies.

#### Neural decoding of hand information in the motor task

We next asked whether hand information could be decoded from each brain region. To this end, we trained a linear population decoder on the neural responses in each brain region (IT/PMv/PFC) during single trials of the left and right hand blocks, and tested the decoder on unseen trials. Decoding accuracy was high in all three brain regions (Figure 3H). The high decoding accuracy in PMv and PFC is concordant with their status as regions that encode motor or task-related information. However, the high hand decoding accuracy in IT is a bit puzzling, and we surmise that it could reflect either motor information, or gaze information arising from monkeys looking at the hand or button before making a movement. Based on the existing analysis, we cannot dissociate these contributions. Nonetheless, we conclude that hand information was reliably present in all three brain regions during the motor task.

### Neural decoding of visual & motor information in the natural task

We next turned to the main purpose of the study, to understand neural responses during real object interactions. Each monkey participated in a session conducted in the large naturalistic arena shown in Figure 2A. We devised a simple natural task that required the monkey to see and interact with specific objects at specific times, so that we could compare neural activity at these times with the neural activity evoked by images of these objects. The natural task is depicted in Figure 4A. On each trial, a human experimenter (either E1 or E2) entered the room with a closed food box, reached the interaction zone, opened the box and offered food (banana, carrot or raisin on separate trials) to the monkey. The monkey had to reach out through the grill to take the offered food from the box with his left or right hand, after which the experimenter left the room (Figure 4A; see Methods for details). During this entire session, we recorded brain activity wirelessly as well as the video from multiple cameras synchronized to the neural data acquisition. We manually annotated the video recorded during each trial with key events that occurred during the trial, for subsequent analyses.

**Figure 4.**
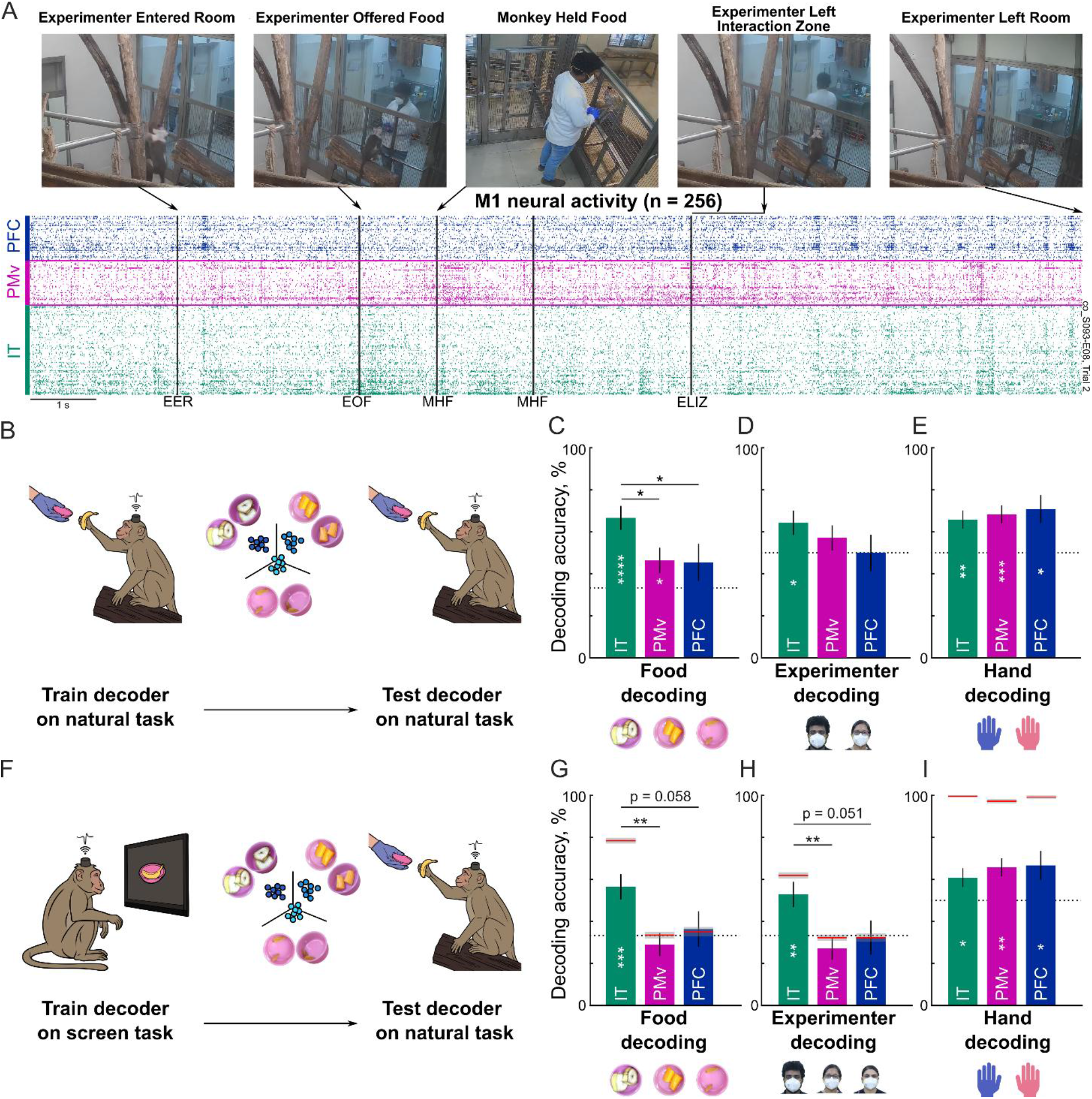
Neural decoding of visual and motor information in the natural task (see also <u>Video S1</u>. (A) Top: Screenshot from an example trial in the natural task, in which a human experimenter (E1) entered the room, offered a banana to monkey M1, and left the room. Bottom: Raster plot of neural activity from all 256 electrodes in monkey M1, with each row depicting the spikes detected in each electrode from PFC (blue), PMv (purple) and IT (green). IT neurons were sorted according to their selectivity for banana images in the touchscreen task, so that the most banana-selective neurons are at the bottom. It can be seen that these banana-selective neurons show bursts of activity particularly around the time of the experimenter offering food (EOF) and monkey held food (MHF) events. Note that there are two MHF events in this trial because the monkey retrieved banana pieces from the box twice, one after the other. Vertical black lines indicate the times of key behavioral events: EER = Experimenter Entered Room, EOF = Experimenter Offered Food, MHF = Monkey Held Food, ELIZ = Experimenter Left Interaction Zone. (B) Schematic of a linear decoder trained on trials in the natural task and tested on unseen trials in the same task. (C) Cross-validated decoding accuracy for the food decoder in IT (*green*), PMv (*purple*) and PFC (*blue*). Error bars represent s.e.m across trials pooled across both monkeys. Asterisks within each bar indicate statistically significant decoding accuracy (* is p < 0.05, **** is p < 0.00005; chi-squared test comparing observed correct/incorrect counts across trials with that expected by chance). Asterisks spanning bars represent the statistical significance of the comparison between brain regions (* is p < 0.05, rank-sum test comparing decoder output across trials). (D) Same as (C) but for experimenter decoding in all three brain regions. *[Face images shown are those of the authors: D.B., S.M. and S.S.]* (E) Same as (C) but for hand decoding in all three brain regions. (F) Schematic of linear decoders trained on the touchscreen task (fixation task for food and experimenter decoding, and motor task for hand decoding) and evaluated on the natural task trials. (G) Cross-validated decoding accuracy for the food decoder in IT (*green*), PMv (*purple*) and PFC (*blue*). All other conventions are as in panel (C). (H) Same as (G) but for experimenter decoding in all three brain regions. *[Face images shown are those of the authors: D.B., S.M. and S.S.]* (I) Same as (G) but for hand decoding in all three brain regions.

The lower panel of Figure 4A depicts the neural activity evoked across all three recorded brain regions in monkey M1 in an example trial in which experimenter E1 offered banana pieces to the monkey. We sorted IT neurons according to their selectivity for banana in the touchscreen task, so that the most banana-selective neurons are at the bottom. It can be seen that neural activity was rich and dynamic, with bursts of activity across the entire neural population, sometimes coordinated across all brain regions, sometimes exclusively within a region, sometimes aligned to specific events, and yet at other times unrelated to the annotated events. Despite this variability, it can also be seen that the banana-selective channels fire strongly around the time of the experimenter offering food (Figure 4A, lower panel, marked as EOF), and the monkey holding food (Figure 4A lower panel, MHF), exactly at the times when the monkey was viewing the banana. This observation suggests that neural activity during the natural task is highly systematic and lawfully related to neural responses to images on the screen.

#### Neural decoding of food & experimenter during the natural task

We next sought to quantify these observations. We first asked whether information regarding food identity (banana/carrot/raisin) or experimenter (E1/E2) could be decoded in the natural task at times when the monkey had to be looking at these items.

To characterize whether food type could be decoded during the natural task, we trained a 3-way linear decoder (Figure 4B) as before on the population activity in each brain region at the time when the monkeys were looking at the food (M1 looked at the food when he was holding it; M2 looked at the food when it was offered; see Figure S1; see <u>Video S1</u>). Cross-validated decoding accuracy for food was significantly above chance in IT and PMv but not in PFC (Figure 4C). While the decoding in IT is presumably due to its encoding of visual features, the decoding in PMv is presumably due to the different grasping movements made by the monkeys to the three food types. This is further corroborated by the fact that food decoding in PMv in the touchscreen task did not differ from chance (Figure 3F).

To characterize whether experimenter identity could be decoded from neural activity in the natural task, we trained a 2-way linear decoder on the population activity in each brain region at the time when the monkeys were looking at the experimenter (for M1, when experimenter offered food; for M2, when experimenter entered room; see Figure S1). Cross-validated decoding accuracy was significantly above chance only in IT but not in PMv or PFC (Figure 4D).

#### Neural decoding of hand used during the natural task

Finally, to characterize whether the hand used by the monkey to receive food could be decoded from neural activity in the natural task, we trained a 2-way linear decoder at times when the monkey was using his left or right hand during the natural task. Cross-validated decoding accuracy was significantly above chance in all three brain regions (Figure 4E). While it is unsurprising that PMv & PFC carry hand information, the availability of hand information in IT could reflect the monkey looking at the hand or food before making a movement.

### Generalization of neural decoding from touchscreen to natural tasks

The above sections show that food, experimenter and hand information can be decoded reliably from neural activity in the touchscreen task and natural task. However, the neural activity patterns corresponding to this information in the natural task may be completely different from the format of information in the fixation task. To investigate this issue, we asked whether decoders trained on neural activity during the touchscreen task could reliably decode the food, experimenter and hand information during the natural task at key events as before (Figure 4F; see Methods). This cross-decoding accuracy was significantly above chance for food and experimenter only in IT (Figure 4G-H), whereas hand information could be reliably decoded from all three brain regions (Figure 4I).

Thus, neural signatures evoked by visual and motor content in the natural task is systematically related to neural activity evoked to these items in the more controlled setting on a touchscreen.

### Neural decoding of face & experimenter viewpoint during the natural task

Since the responses of IT neurons are not perfectly viewpoint invariant, we wondered whether neural responses during the natural task would reflect the viewpoint at which objects were viewed in the natural task. To this end, we trained separate food and experimenter decoders for each viewpoint of these images in the fixation task, and tested their accuracy during the natural task. We found that decoders trained on the oblique views of the food box and on the profile view of the experimenters had the highest accuracy (Figure S2). These are concordant with our observations during the experiment, that the monkeys viewed the food box in an oblique view as they took the food from the box and viewed the experimenters from the side view as they entered the room.

### Neural correlates of food anticipation

Although three food types were offered to the monkeys during the natural task, we wondered whether the monkeys were anticipating any particular type of food on each trial. We reasoned that, if the monkeys were anticipating a banana in a given trial, the food decoder trained on the fixation task would show a high banana probability even before the experimenter offered food. Indeed, we observed signs of food anticipation by both monkeys (Figure S3; see also <u>Video S1</u>).

### Variability and fidelity of neural signatures in natural & screen tasks

The above sections show that food, experimenter and hand information can be reliably decoded from the screen task as well as the natural task. Furthermore, neural signatures are similar in the two tasks, given the successful cross-decoding from one task to the other. However, these analyses do not directly compare the variability of neural responses or the fidelity of object information in the two tasks. By fidelity we mean the response variability present along the dimensions relevant to decoding object information. We specifically evaluated two possibilities: (1) IT neurons might show greater variability in the natural task, both along object-relevant and irrelevant dimensions; (2) IT neurons might show greater variability in the natural task, but only along object-irrelevant dimensions.

#### Variability of neural signatures in natural ḥand screen tasks

We compared the overall level and variability of neural activity in the natural and screen tasks. In the screen tasks, we considered the spikes evoked in a window 0-0.35 s relative to image onset in the fixation task and similar window relative to movement onset in the motor task (see Methods). In the natural task, we considered the spikes evoked in an identical duration window centered around each key event in the task.

Firing rates of IT visual neurons in the visual task were higher in the fixation task, compared to nearly all events in the natural task, except for when the experimenter offered food (Figure 5A). This is presumably because the visual task involves sudden image onsets from a black background whereas the natural task involves more gradual visual changes. IT neurons also had higher firing rates in the fixation task compared to the motor task (Figure 5A). This is expected since the fixation task involved diverse natural images whereas the motor task involved buttons of the same color.

**Figure 5.**
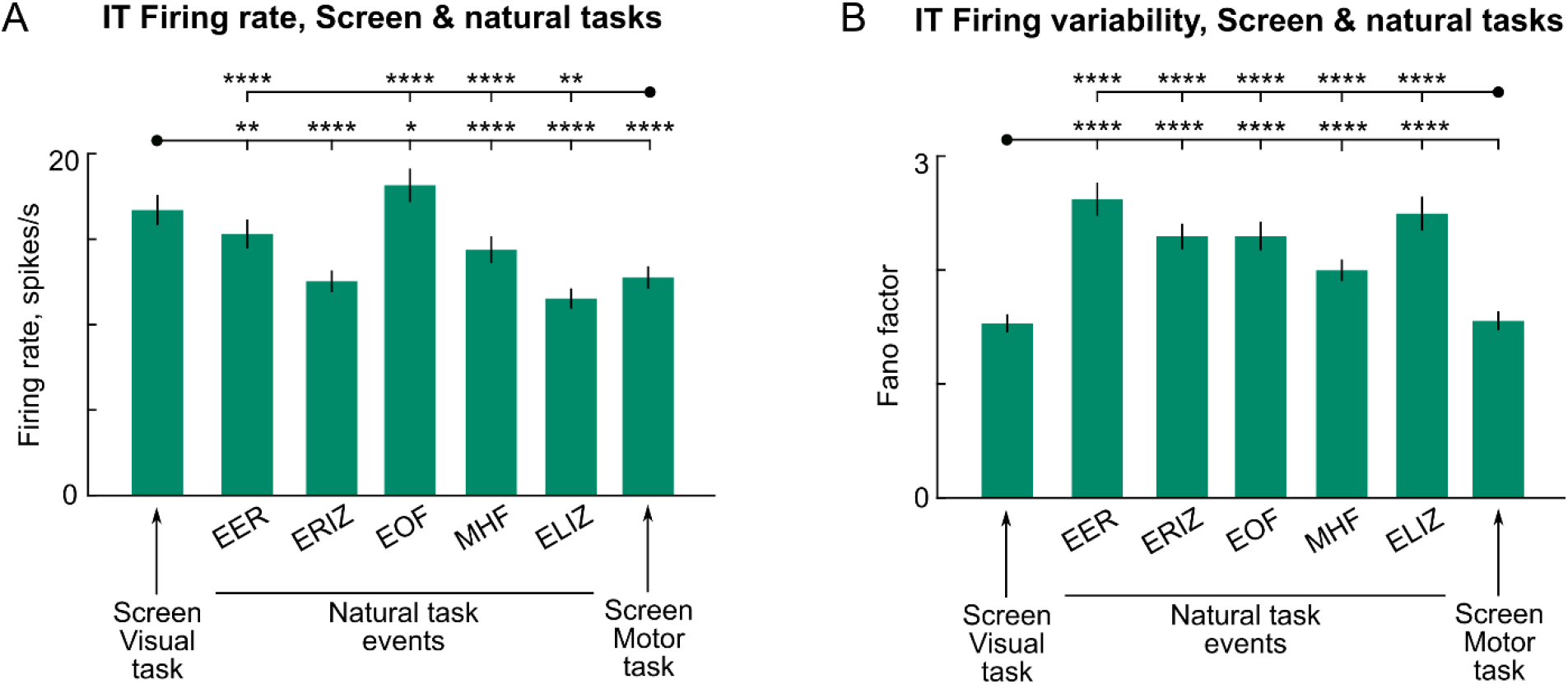
Neural variability during natural and screen tasks. (A) Average firing rate of IT neurons during screen & natural tasks in equivalent time windows, with error bars indicating s.e.m across neurons. Asterisks indicate statistical significance of the corresponding comparison (* is p < 0.05, ** is p < 0.005, etc; sign-rank test across firing rates of 190 visual neurons; comparisons shown only between each screen task marked by a dot with all the key events in the natural task). It can be seen that neural responses were higher for the screen task compared to all natural task events except for when the experimenter offered food (EOF). Event abbreviations: EER = Experimenter Entered Room, ERIZ = Experimenter Reached Interaction Zone, EOF = Experimenter Offered Food, MHF = Monkey Held Food, ELIZ = Experimenter Left Interaction Zone. (B) Average Fano factor of IT neurons during screen & natural tasks in equivalent time windows, with error bars indicating s.e.m across neurons. For each neuron, the Fano factor was calculated as the variance/mean of spike counts across trials of a given condition in each task and then averaged across all the conditions. Asterisks indicate statistical significance as in panel (A). Note that firing rate variability was generally higher for all natural task events compared to the screen tasks.

To compare variability of neural activity in the natural and screen tasks, we calculated the Fano factor, a commonly used measure of neural variability. The Fano factor is 1 for a Poisson process independent of its rate, and cortical neurons are usually observed to have a Fano factor slightly larger than 1 (Vogels et al., 1989; Baddeley et al., 1997; Shadlen and Newsome, 1998; McAdams and Maunsell, 1999; Sripati and Johnson, 2006). For each neuron, we calculated the variance divided by the mean of the spike count across trials of each unique condition and averaged it across conditions. Fano factor for the visual task was significantly lower on average compared to all other events in the natural task (Figure 5B). The greater Fano factor during the natural task events is expected given the many additional variables encoded in the brain during natural movements.

In sum, we find that IT neurons have lower firing rate and greater variability during the natural task compared to the screen task. We found similar results for motor neurons in PMv & PFC (Figure S4).

#### Food decoding fidelity in the natural & screen tasks

The above analysis shows that IT neurons show greater variability in the natural task. However, this variability may or may not affect the fidelity of food information. We started by comparing the food decoding accuracy in the screen task and natural task (78.5% in the screen task vs 66.7% in the natural task; Figure 3F vs Figure 4C). However, this is not a balanced comparison, since each decoder was trained on different numbers of trials (∼400 trials in the fixation task vs only ∼30 trials in the natural task), and any difference in accuracy could be due to the difference in training data.

We therefore trained decoders on exactly equal numbers of trials in the screen task (taking equal numbers of trials for each food view) and in the natural task (see Methods). We found that the food decoder accuracy was higher in natural task, but this difference did not attain statistical significance (Figure 6A).

**Figure 6.**
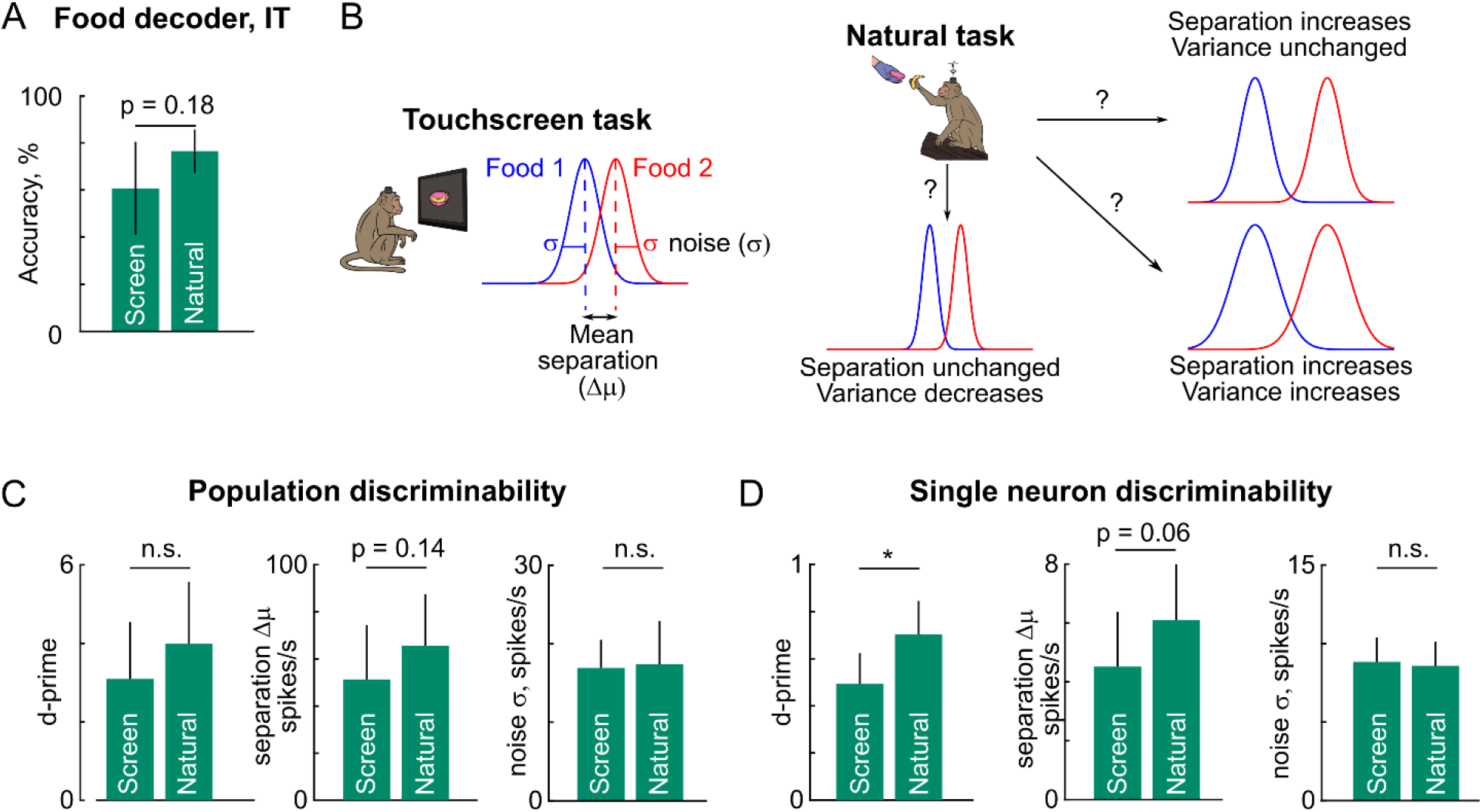
Fidelity of object information during natural and screen tasks. (A) Schematic showing the distribution of projections of population activity along the linear classifier vector for Food 1 (*blue*) vs Food 2 (*red*). To evaluate the fidelity of information in each task, we compared the mean separation between the classes (Δ*μ*), the noise within each class (*σ*) and the overall discriminability (*d*^′^ = Δ*μ*/*σ*). (B) Schematic showing three possible explanations for increased decoding accuracy in the natural task compared to the fixation task: either the separation increases without change in noise, separation is unchanged with a decrease in noise, or both separation and noise increase. (C) Population discriminability, separation and noise levels for IT neurons, with error bars representing standard deviation across bootstrap samples. For details of each calculation, see Methods. Asterisks represent statistical significance, calculated as the frequency of violations in the depicted trend across bootstrap samples (n.s. is p > 0.05, * is p < 0.05). Note that discriminability is slightly larger in the natural task, driven by an increased separation without any increase in noise. (D) Same as (C) but using single-neuron analyses. For each neuron we calculated the discriminability, separation and noise levels for separating classes (see Methods). Asterisks represent statistical significance as before. Note that single neuron discriminability is larger in the natural task, and this is driven by an increased separation without any increase in noise levels.

**Figure 6.**
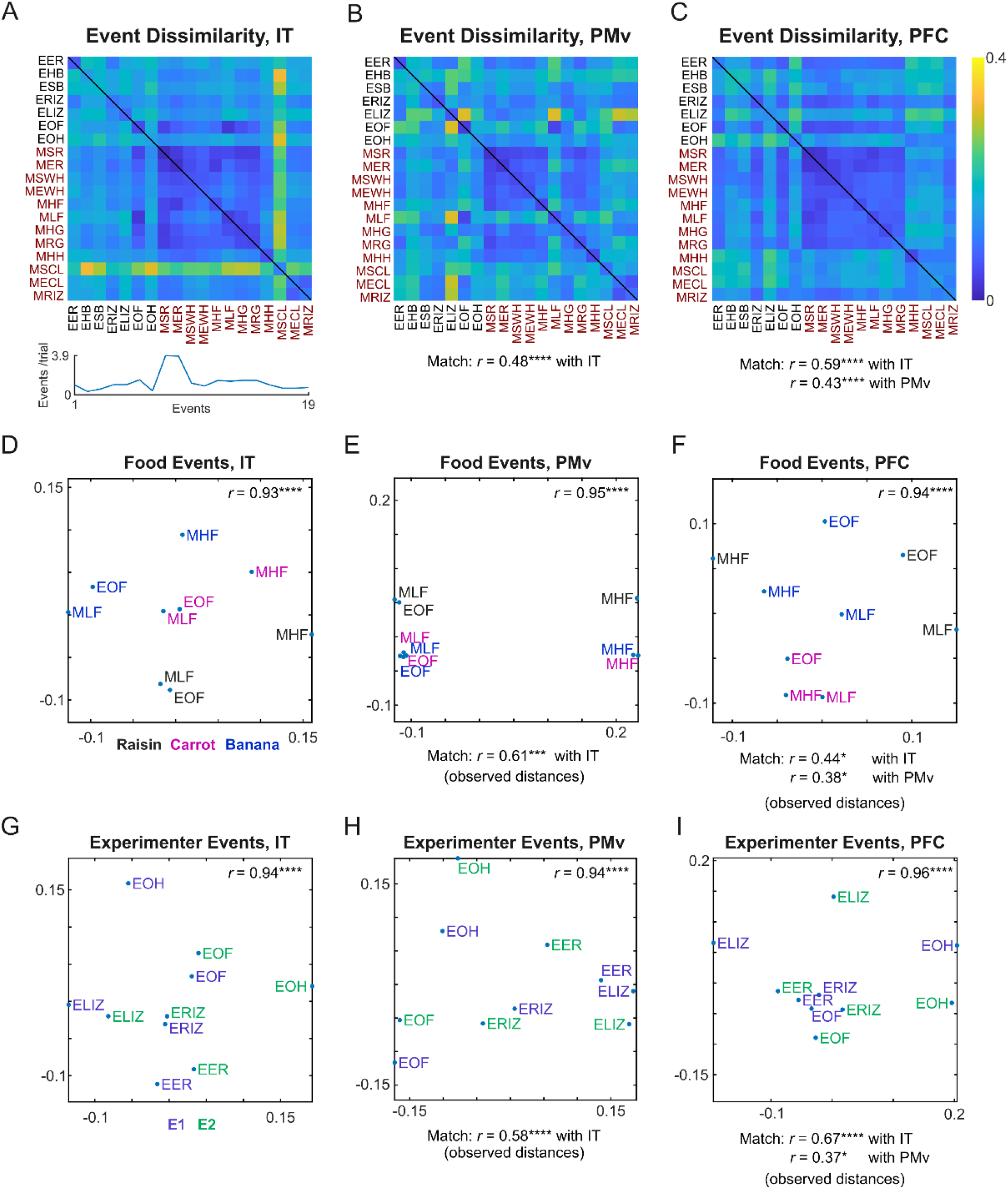
Neural dissimilarity between key behavioral events in the natural task. (A) *Top:* Neural dissimilarity in IT between all pairs of annotated events in the natural task, calculated as the correlation distance (1 – r) between the population vector of firing rates in a window −0.25 s to 0.25 s relative to each event, averaged across all event instances in a given monkey, which is then averaged across monkeys. Values along the main diagonal represent the correlation distance between two halves of the instances of each event, which should be ideally zero if the neural firing rates are perfectly repeatable. *Bottom:* Average number of occurrences per trial of each event pooled across all trials in both monkeys. Event abbreviations: EER, Experimenter Entered Room; ESB, Experimenter Show Box; EHB, Experimenter Hide Box; ERIZ, Experimenter Reached Interaction Zone, ELIZ, Experimenter Left Interaction Zone; EOF, Experimenter Offered Food; EOH, Experimenter Offered Hand; MER-Monkey End Reach; MSR, Monkey Start Reach; MSWH, Monkey Start Withdraw Hand, MEWH, Monkey Stop Withdraw Hand; MHF, Monkey Held Food, MLF, Monkey Look At Food; MHG, Monkey Held Grill; MRG, Monkey Released Grill; MHH, Monkey Held Hand; MSCL, Monkey Start Climb Log; MECL, Monkey End Climb Log; MRIZ, Monkey Reached Interaction Zone. (B) Same as (A) but from PMv. The correlation below indicates the match between PMv event dissimilarities with that of IT (across ^19^C_2_ pairs = 171 dissimilarities). (C) Same as (A) but from PFC. The correlations below indicate the match with IT & PMv. (D) Multidimensional scaling plot of event dissimilarities in IT for a subset of events that involved food, but colored separately for by food (*blue*, banana; *magenta*, carrot; *black*, raisin). In the plot, events are nearby if they elicited similar neural activity. The correlation at the top shows the match between pairwise event dissimilarities and the 2D distances in the plot (asterisks represent statistical significance: **** is p < 0.00005). (E) Same as (D) but for PMv. The correlations below the plot show the match between pairwise event dissimilarities in PMv with those of IT. (F) Same as (D) but for PFC. The correlations below show the match with IT & PMv. (G) Multidimensional scaling plot of event dissimilarities in IT for a subset of events that involved the human experimenter, but colored separately by experimenter (*blue*, E1; *green*, E2). (H) Same as (G) but for PMv. (I) Same as (H) but for PFC.

We reasoned that the slight advantage of food decoding in the natural task could arise from three possibilities (Figure 6B). Compared to the fixation task, the natural task could have better separation between the food types with the same noise, better separation with increased noise, or same separation but smaller noise (Figure 6B).

To evaluate these possibilities, we trained a linear decoder on each pair of food items and projected the population activity of each trial along the classifier vector. We then calculated the average separation between classes (Δ*μ*), the average noise level within each class (*σ*), and a discriminability (d’) measure which is simply the ratio of the two (*d*^′^ = Δ*μ*/*σ*). At the population level, we found that the population discriminability and mean separation were slightly higher in the natural task compared to the screen task, but this difference was not statistically significant in either case (Figure 6C). By contrast, the noise level was not significantly different in the natural task.

To be sure that the results were not due to high-dimensional classifiers involved, we performed a similar analysis at the single neuron level. For each neuron and food pair, we calculated the average difference in firing rate for the two classes (Δ*μ*), the average standard deviation of firing rate within each class (*σ*), and the discriminability (*d*^′^ = Δ*μ*/*σ*), as detailed in the Methods. We found that the discriminability in the natural task was significantly higher, and this effect was driven by an increased separation between the classes (Figure 6D).

We conclude that the fidelity of food decoding was slightly better in the natural task despite the greater variability overall of neural firing. We obtained similar results for experimenter decoding (Figure S5).

Taken together, these findings show that brain activity during real object interactions is strikingly reliable and comparable in fidelity to brain activity while viewing images. This is striking considering that natural behaviors involve large variations in sensory inputs and motor outputs across trials.

### Neural signatures of key behavioral events in the natural task

Since the natural task consisted of key behavioral events, we wondered if each of these events evokes a unique neural signature in the recorded brain regions. To investigate this possibility, we took the average firing rate evoked across the entire population in a window centered around each key event, and asked how similar this firing rate was across every other event (see Methods).

The resulting event dissimilarity matrix is shown in Figure 7. It can be seen that event dissimilarities were smaller for all movement-related events (brown-colored events in Figure 6A-C). Neural dissimilarities between all regions were significantly correlated (*r* = 0.48 for IT-PMv; *r* = 0.59 for IT-PFC; *r* = 0.43 for PMv-PFC; p < 0.00005). Thus, neural signatures of key events were similar across all regions. These event dissimilarities were similar within each monkey as well (Figure S6-7).

Since IT encodes visual information, we hypothesized that events involving each food type or each experimenter would cluster together. Likewise, since PMv encodes motor information, we expected that events involving similar movements would cluster together. To evaluate this possibility, we performed a multidimensional scaling analysis of food-related events after separating them by food. It can be seen that food-related events revealed clustered by food type in IT (Figure 6D), by action in PMv (Figure 6E) and by food type in PFC (Figure 6F). Similarly, a multidimensional scaling analysis of experimenter-related events revealed that actions clustered together regardless of experimenter identity in all three brain regions (Figure 6G-H-I).

Thus, key events in the natural task elicit distinct neural signatures that reflect event similarity as well as the encoding priorities of each brain region.

## DISCUSSION

Here, we set out to investigate how neural responses evoked during natural interactions with real objects are related to neural responses evoked by images of these objects. To this end, we performed wireless recordings in high-level sensory and motor regions (IT, PMv, PFC) of freely moving monkeys performing a natural task in a large natural arena and also performing controlled tasks on a touchscreen workstation. We report three main findings: (1) neural activity during real object interactions bears a lawful relationship with neural responses to images of these objects; (2) neural activity during real object interactions was weaker and more variable than to images, but the variability along object-relevant dimensions was strikingly similar; and (3) key events during the natural task elicited reliable neural signatures while also reflecting the encoding priorities of each brain region. These findings represent an important first step in understanding the neural basis of real-world vision. Below we discuss each of these findings in context of the existing literature.

### Relationship between real objects and images

We have found that neural activity in IT neurons during interactions with real objects is lawfully related to neural activity evoked to images of these objects. We demonstrate this by showing that neural activity decoders trained on images of food and faces can accurately identify the food and experimenter identity during the natural task. Our finding is concordant with many studies of IT neurons showing similar responses while viewing 3D objects and their images (Tanaka et al., 1991; Kobatake and Tanaka, 1994; Connor et al., 2007; Orban, 2011). However, these results were obtained while animals viewed these objects in the absence of any interactions. By contrast, during natural interactions, real objects contain rich stereo, motion and parallax cues, might be seen from differing viewpoints, subject to multiple eye movements and finally could be manipulated differently each time during interactions (Snow and Culham, 2021; Culham and Deligiannis, 2026). Our findings assume greater importance by showing that during natural behaviors, despite the inherent variability, IT neurons exhibit similar neural signatures during interactions with real objects and their images.

The accurate decoding of real objects, however, was possible only at certain key events in the natural task, where we were reasonably certain that the monkeys were looking at these objects. This was confirmed by our own observations as well as by the decoding accuracy. For instance, food decoding was most accurate for Monkey M1 when he was holding the food, but was most accurate in Monkey M2 when the experimenter offered food (Figure S1). Thus, neural responses during real-world interactions with objects will need to be interpreted in light of the monkeys’ behavior.

### Variability of neural responses during real object interactions

Natural behaviors are complex and variable: even the simple action of eating food could involve seeing the food from different viewpoints each time and reaching differently each time. The prevailing belief is that neural activity during such natural tasks will be highly variable and hard to decode (Miller et al., 2022; Cisek and Green, 2024; Leopold, 2024). By contrast, if the brain is indeed optimized for natural behaviors, this additional variability should not affect the encoding of relevant information (Miller et al., 2022; Parodi et al., 2025). Our results confirm this latter possibility, by showing that, even though neural activity is indeed more variable in the natural task, food information is encoded with striking fidelity, similar or even better than images shown on the screen (Figure 6). This is similar to the observation in motor cortex that neural activity related to motor planning occurs in a null space to avoid interference with ongoing movement execution (Churchland and Shenoy, 2024).

### Reliability of neural signatures during natural behaviors

We have found that key events in the natural task are associated with reliable neural signatures. This is consistent with previous studies that have shown encoding of navigational variables (Maisson et al., 2023) and complex actions (Voloh et al., 2023; Lanzarini et al., 2025) in higher order motor and prefrontal regions during natural behaviors. However, we also find that key events in the natural task are encoded slightly differently in IT and PMv. Events clustered by food and experimenter in IT, whereas they clustered by motor actions in PMv, consistent with their known status as visual and motor regions. Thus, key events in the natural task are not only associated with distinct neural signatures, but also reflect the encoding priorities of each brain region.

### Limitations of our study

Our study represents a first step towards understanding the neural basis of real world vision by comparing neural activity during real object interactions with neural responses to images on a touchscreen. However, our study has several limitations. First, although we have demonstrated an interpretable path between neural responses to images and neural activity during natural behavior, we did not have complete information about the visual input seen by our animals in the natural setup, nor did we know their gaze. These will require head-mounted cameras and eye tracking, which are beyond the scope of our present study. There have been exciting developments in this direction in marmosets (Singh et al., 2025), and extending them to larger primates will be interesting possibilities for future work. Second, we did not have 3-dimensional joint positions of our animals in the natural arena, which limited our analysis of their movements. These will require multi-camera setups like those reported recently (Maisson et al., 2023; Voloh et al., 2023). In general, an increasing number of signals and variables will be critical to interpreting and understanding brain activity during natural behaviors, along with recording brain activity also in more controlled settings.

### Conclusions

Taken together, our results show that brain activity during natural behaviors can be understood by comparing it with neural responses in more controlled settings. We propose that understanding natural behavior will require studying brain activity in well-controlled settings where a variety of signals can be recorded as well as in fully natural settings where not every signal can be recorded (Cisek and Green, 2024; Parodi et al., 2025).

## METHODS

All animal experiments were performed according to a protocol approved by the Institutional Animal Ethics Committee of the Indian Institute of Science, Bangalore (CAF/Ethics/750/2020) and by the Committee for the Purpose of Control and Supervision of Experiments on Animals, Government of India (V-11011(3)/15/2020-CPCSEA-DADF). Many experimental procedures are common to previously reported studies from our laboratory (Ratan Murty and Arun, 2015; Jacob et al., 2021), and summarized briefly below.

### Neurophysiology

We implanted two adult bonnet macaque monkeys (Ma*caca radiata,* M1: *Co* & M2: *Di*, both aged 9.5 years) with Floating Microelectrode Arrays (FMA) from Microprobes for Life Science using standard neurophysiological procedures reported previously (Ratan Murty and Arun, 2015), and summarized here along with novel methodologies unique to these recordings. Surgeries were planned using CT & structural MRI scans on both monkeys, co-registered using 3D Slicer. Array locations were verified during surgery based on sulcal landmarks visible through the craniotomy, and their precise locations were estimated by aligning photographs of the sulcal landmarks taken during surgery with the sulci visible from the structural MRI (Figure 1C).

Each monkey was implanted with 8 FMAs of 32 channels, resulting in a total of 256 electrodes. Each FMA had 32 single-shank microelectrodes arranged in a 4 x 8 grid with an inter-electrode spacing of 400 microns. Monkey M1 was implanted with 4 FMAs into the central portion of the left inferior temporal (IT) cortex, 2 FMAs in the left ventrolateral prefrontal cortex (PFC), and 2 FMAs in the left ventral premotor cortex (PMv). Monkey M2 was implanted with 4 FMAs in the right inferior temporal cortex and 4 FMAs in the right ventral premotor cortex. In Monkey M1, the array centers ranged from 2.4-6.4 mm anterior to the AP0 (ear bars) for the IT arrays, and AP 27.0-31.8 mm for PMv & PFC arrays. In Monkey M2, the array centers ranged from AP 2.7-10.2 mm for IT arrays and AP 22.5-27.2 mm for PMv arrays.

In both monkeys, each pair of wires from electrode arrays were passed through a custom-designed titanium pedestal and connected to 64-channel headstage. The pedestal was fixed to the skull along with an H-plate and cemented with dental acrylic. The electrode array cable entry into the pedestal was made liquid-proof during surgery using epoxy glue (Kwik-Sil, World Precision Instruments, Inc). In all, there were four 64-channel headstages and one wireless interface boards (White-Matter LLC) in a vertical stack along with a retainer – these remained inside the pedestal for the entire study. To protect the electronics inside the pedestal, we designed a 3D printed nightcap with the same dimensions as the wireless logger, which was fixed to the pedestal using screws. To make its compartment liquid-proof, we devised a slot in the pedestal into which we inserted a rubber O-ring that would make a tight seal with the nightcap or wireless logger and prevent any liquid ingress.

### Wireless recordings

Monkeys were trained to enter a custom-designed monkey chair, and were snout restrained for around 10 −15 minutes before implantation (Jacob et al., 2021). On the day of recording, each monkey was brought into the chair, and its snout was restrained temporarily for about 5 minutes to clean its implant, remove the nightcap and fix the wireless logger to the titanium pedestal using screws. Extracellular wideband signals were logged at 25 KHz using a custom wireless logger that was synchronized to a neural data acquisition system (*eCube,* White-Matter LLC). These wideband signals were high-pass filtered and thresholded using a commercial software (*OfflineSorter*, Plexon Inc) to obtain multiunit activity (cutoff frequency 250 Hz, Butterworth 4-pole filter; threshold −3.75 sigma, waveform length 1.6 ms, pre-threshold period 0.32 ms and deadtime 1.28 ms).

### Touchscreen workstation

Monkeys performed the screen-based tasks on a touchscreen workstation with accurate eye-tracking capabilities as reported previously (Jacob et al., 2021). The workstation was under control of the software NIMH MonkeyLogic version 2.2.1 (Hwang et al., 2019), and eye tracking was performed at 120 Hz (ISCAN Inc, ETL 300HD). In addition, the monkeys movements were monitored from three viewpoints (front, top and side) using video cameras that were synchronized to the neural data acquisition system (e3Vision, WhiteMatter LLC, 1280 x 720 resolution, 30 fps). To track possible timing differences between the issuing of display commands and the actual appearance of visual stimuli on the screen, we also presented a white square at the top of the touchscreen along with each stimulus, which provided onset input to a photodiode (TSL-257). The white square and photodiode were covered with an opaque acrylic cover so that they were not visible to the monkey. The output of the photodiode was recorded at 25 KHz by the neural data acquisition system, and was used to correct all visual event appearance times.

### Alignment of video recordings to neural data

Although video cameras at the touchscreen workstation were fully synchronized with the neural data acquisition system, the video cameras in the natural arena were not synchronized in this manner. To achieve complete alignment, we noted that some of the synchronized video cameras as well as some non-synchronized cameras had a view of the touchscreen. We therefore created a synchronization task, whereby full screen white images flashed at precise times on the touchscreen, which were then captured by the photodiode (to get precise onset times) and also by all the cameras. We used the rise and fall of image intensity to align the timing of all cameras to the neural data acquisition, and verified them by comparing the timing of events shown in all cameras.

### Natural task

To study natural vision, we devised a simple behavioral task. On each trial, a human experimenter entered the natural arena with a covered box of food, approached the interaction zone and opened the lid of the box to reveal the food, upon which the monkey took the food from the box. In all, one of three types of food (pieces of banana, carrot or raisin) were offered by one of two experimenters (the authors D.B & S.M). The food box was offered so that it was close to the monkeys’ right (dominant) hand in the first 24 trials and to the left hand in the next 12 trials. The sequence of trials was randomly chosen using a computer with the constraint that consecutive trials would not have the same food or experimenter. There were 37 trials for monkey M1 and 36 trials for monkey M2.

We manually annotated the videos which contained the clearest view of the monkey-human interaction for the times of key events during each trial. The key events were chosen after carefully reviewing the entire session video to identify distinct, repeating and well-defined events that occurred across trials. The annotated events were: Experimenter Entered Room (EER), Experimenter Left Room (ELR), Experimenter Show Box (ESB), Experimenter Hide Box (EHB), Experimenter Reached Interaction Zone (ERIZ), Experimenter Left Interaction Zone (ELIZ), Experimenter Offered Food (EOF), Experimenter Offered Hand (EOH), Experimenter Withdrew Hand (EWH), Monkey Start Reach (MSR), Monkey End Reach (MER), Monkey Start Withdrew Hand (MSWH), Monkey Stop Withdrew Hand (MEWH), Monkey Held Food (MHF), Monkey Held Hand (MHH), Monkey Held Grill (MHG), Monkey Released Grill (MRG), Monkey Face Experimenter Entered Room (MFEER), Monkey Look At Food (MLF), Monkey Look At Empty Box (MLEB), Monkey Look At Experimenter (MLE), Monkey Look At Experimenter Hand (MLEH), Monkey Start Climb Log (MSCL), Monkey End Climb Log (MECL), Monkey Reached Interaction Zone (MRIZ), Monkey Left Interaction Zone (MLIZ), Monkey Startled (MS).

### Touchscreen tasks

Before the natural task recordings, both monkeys performed two touchscreen-based tasks: a visual task and a motor task.

#### Eye calibration block

Each task started with a block for calibrating eye position. The monkey had to initiate a trial by touching a circular hold button (radius 4°, 28° away from the center) within 10 seconds. Once the trial was initiated, four gray circles (radius 0.5°) appeared consecutively at four corners of a square of side 16° in random order. The monkey had to look within 10° (for M1) or 8° (for M2) of each of these circles within 0.5 s, for 0.3 s, as each appeared on the screen, and received a juice reward for fixating all 4 targets successfully. The trial was aborted if the monkey failed to fixate on any of the 4 target circles. The monkey had to complete two correct calibration trials. The eye data collected from these trials was used to learn a linear transformation between the eye signal values (x & y separately) at each known gray circle position, so that new (x, y) eye signal values could be transformed to obtain the position of the eyes in degrees of visual angle.

### Visual task

Each monkey performed a fixation task to characterize visual responses. Each trial started with a blue circle (radius 4°, presented 28° away from the center) which the monkey had to touch within 10 s. Because of individual differences in touching behavior, we adjusted the tolerance around the hold button so that M1 had to touch within 8° of the hold button whereas M2 had to touch within 2° of the hold button. After the monkey touched hold button, a yellow dot (radius 0.1°) appeared at the center on which the monkey had to maintain gaze within a 10° radius within 0.5 s of its appearance. After maintaining fixation for 0.3 s, a sequence of 8 images was presented for 0.2 s each with a 0.2 s blank interval, during which the monkey had to maintain fixation within this radius and received a juice reward. The trial was aborted if the monkey failed to maintain hold or broke fixation, and such error trials were repeated after a random number of other trials. Although we used a fairly liberal fixation window because we were unsure about fixation quality, our post-hoc analyses of the data revealed that monkeys maintained accurate fixation throughout the trial (standard deviation of eye position during each trial: 1.05° for horizontal and 0.99° for vertical for M1, and 0.71° for horizontal and 0.91° for vertical for M2, averaged across trials; mean jitter in eye position across trials: 0.37° for horizontal and 1.3° for vertical for M1, and 0.58° for horizontal and 0.80° for vertical for M2).

In the main fixation task, the monkey saw three groups of images in separate blocks: (1) images of the food box that was either empty or containing pieces of banana, carrot or raisin, matched to the likely views of food seen by the monkeys during the natural task (3 food types x 5 views in the box = 15 images, 5 images of empty box from 5 views, 3 food types x 3 views of these pieces on the floor = 9 images, 3 images of empty floor; total 32 images); (2) images of the human experimenters (either E1 or E2 who would subsequently participate in the natural task, or E3 who served as a control; 3 experimenters x 2 variations with/without face mask x 5 views = 30 images) and (3) 18 images sampled from the natural arena. Each category of images was shown scaled in size so that they matched roughly the size at which they were likely to be viewed in the natural task (food images: 7.8° along the longest dimension; face images: 22.2°; play area images: 40.8° along the longest dimension). In all, there were 80 unique images in the experiment, with each repeated 16 times over the course of 160 correct trials. Images within a trial were shown in random order with the constraint that consecutive images do not contain the same viewpoint, food type or face in the food and face blocks, and were randomly interleaved in the natural arena image block such that two consecutive images will not be from same category (e.g. perch or log categories).

All data analysis in the fixation task were performed on a window 0 – 0.3 s after image onset for food images and 0-0.3 s after image onset for experimenter faces. Varying these choices yielded qualitatively similar results.

### Motor task

Each monkey performed a center-out reach task with his left and right hand in separate blocks. We ensured that the monkey would use a particular hand by affixing a grill to the opposite side of the juice spout so that only that hand would be convenient to use. Each task started with a gray button (radius 4°, presented 15° from the center) appearing on the screen which the monkey had to touch to initiate the trial. Upon maintaining hold for 0.3 s, the button disappeared and a green circle (radius 4°) appeared at one of four locations (up/down/left/right, 13° away). The monkey had to touch this button within 5 s to complete the trial and receive juice reward. Trials corresponding to each reach location were presented 16 times each, for a total of 64 trials. Error trials were repeated after a random number of other trials. Because of individual differences in touching behavior, we adjusted the tolerance around the hold button so that M1 had to touch within 8° of the hold button whereas M2 had to touch within 2° of the hold button. All data analyses were performed on a window −0.5 s to 0.5 s relative to the time when the monkey touched the target. This time window was chosen so that it captured most phasic neural responses that were observed time locked to movement onset.

### Sorting neurons by banana selectivity (Figure 4A)

To calculate banana selectivity, we calculated the average firing rate (window of 0 s - 0.3 s from target image onset) for each neuron to all banana images with food box and likewise for carrot and raisin responses. We then calculated the difference between the average banana response and the average of the carrot and raisin responses, and sorted cells in descending order of this selectivity for visualization.

### Linear decoders for food, experimenter and hand decoding (Figure 3-4)

To evaluate the amount of information present in neural activity, we performed a linear decoding analysis. In the case of food decoding in the touchscreen visual task, we took the firing rate (in the time window specified above) on each trial of a given image, and created a vector of firing rates by pooling across all neurons of a given brain region. This resulted in a collection of N x M population activity vectors (corresponding to N correct trials and M neurons) along with the corresponding food label (banana/carrot/raisin) for each trial. To remove redundancy in these population vectors, we performed a dimensionality reduction using principal component analysis (PCA) – this resulted in a new set of population vectors that captured a certain percentage (80% for hand decoding, 90% for food decoding & 99% for face decoding) of the firing rate variance. To evaluate this linear decoder on the touchscreen task (Figure 3), we performed a 5-fold cross-validation, whereby the decoder was trained on 80% of the data (*classify* function in MATLAB 2025) and tested on the held-out 20%, and by concatenating the performance on the held-out splits, we were able to obtain the decoding accuracy for all trials where the decoder had never seen these trials during training.

To evaluate this decoder on the natural task (Figure 4G), we calculated the firing rates from each neuron in 0.02 s bins across a time window of 0 s to 0.3 s relative to the event onset (for M1, this was when monkey held food; for M2, this was when experimenter offered food). To calculate decoder output, we subtracted the mean firing rate of the trial (since that is done during PCA) and then projected the population activity vectors along the same principal components as before, and passed the resulting vectors to the decoder to obtain predicted labels for each 0.02 s bin. For each trial, we took the most frequent decoder label across all bins in the time window, on the presumption that this would capture the possibly variable time at which the monkey might have looked at the food. We obtained qualitatively similar results upon taking the decoder label from the mean activity across the entire window, and the mean label across all bins instead of the most frequent label.

For experimenter decoding, we proceeded likewise except that the labels were experimenter identity (E1/E2/E3) and pooled across faces with and without masks and we tested these decoders in the natural task for the event “Experimenter Offered Food” for M1 and “Experimenter Entered Room” for M2. We calculated the firing rates from each neuron in 0.02 s bins for monkey M1 and monkey M2 respectively, across a time window of 0 s to 0.3 s relative to the event onset.

For hand decoding, the labels were the hand used during the motor task (left/right), and the decoder was tested in the natural task for events when the monkey used his left or right hand (regardless of whether the other hand was being used). We calculated the firing rates from each neuron in 0.02 s bins across a time window of −0.5 s to 0.5 s relative to the event onset.

The decoders were also evaluated within the natural task (Figure 4C,4D,4E). We first calculated the PSTH (binsize = 0.02 s ms for both monkeys) for all the trials. Then we averaged neural activity for bins which fell within a window of the event of interest. Then, we train a leave-one-out linear classifier to classify hand used, food and face seen by the monkey during the natural task. First, we did PCA to remove the redundancy on the training data and then we trained the classifiers on all but one trial from natural task. Then we subtracted the mean firing rate of trial from the left-out data and projected it into the PC space of the training set. Then we used *classify* function from MATLAB 2022b to obtain the labels for food, face and hand decoders. The parameters remain same as the previous screen to natural task decoders.

### Analysis of Neural Variability (Figure 5)

To compare the variability of the neural activity across tasks we calculated the firing rates for the visual task, events in the natural task and the motor task in identical time windows of duration 0.35 s. We obtained qualitatively similar results upon varying this choice. The analysis window was adjusted for each task to capture the major modulation in firing rate. These windows were 0 s to 0.35 s after image onset for the visual task, −0.4 s to −0.05 s for the right-hand task (relative to movement onset), −0.05 s to 0.3 s for the left-hand task (relative to movement onset) and 0 s to 0.35 s from the event of interest in the natural task. We selected five key events from the natural task for the analyses: Experimenter Entered Room (EER), Experimenter Reached the Interaction Zone (ERIZ), Experimenter Offered Food (EOF), Monkey Held Food (MHF), and Experimenter Left Interaction Zone (ELIZ).

To calculate the firing rates for the visual and motor tasks we obtained the average firing rates for each condition across trials and then averaged the firing rates across neurons in both monkeys. To calculate the average firing rates for the natural task, we aligned the spikes to the event of interest in the natural task and then calculated the average firing rates across conditions and repetitions, and then averaged across neurons in both monkeys.

To calculate the Fano factor for the visual and the motor task, first we obtained the spike counts for each condition – 24 conditions (3 food x 8 views) for the visual task and 8 conditions (2 hands x 4 reach directions) for the motor task. For each condition we calculated the ratio of the variance of the spike counts and their mean. Then, we averaged the Fano factor across conditions and then averaged it across all visual neurons. To calculate the Fano-factor for the natural task, we obtained spike counts in window of 0 s-0.35 s around event of interest in the natural task. Then using the method above we calculated the Fano-factor for each condition in natural task (6 unique conditions: 3 food types x 2 experimenters). Then, we averaged the Fano-factor across these six conditions and then finally took an average across the visual neurons.

### Analysis of neural discriminability (Figure 6)

To compare the fidelity of food information in the screen task and natural task, we performed an additional analyses on the population decoders trained on food in the two tasks. Since the differing numbers of trials in the two tasks can lead to differences in decoding accuracy and discriminability, we performed a matched decoding analysis, as detailed below.

#### Neural discriminability in the fixation task

Since there were fewer trials in the natural task (n = 36; 3 food x 12 trials), we selected a matched number of trials from the fixation task. To avoid biased sampling, we selected equal numbers of trials of each food type as well as equal numbers of trials of each food view. We then trained a linear classifier to decode the food type from the fixation task and calculated its cross-validated decoding accuracy using leave-one-out cross-validation. For each pair of food items, the linear classifier returns a classifier vector that best separates these two classes. To calculate the population discriminability, we projected the neural activity vector corresponding to each trial along the classifier vector, which returns a scalar projection for each trial. The average separation between classes (Δ*μ*) was then simply the difference between the average projections of the two classes. The noise (*σ*) was calculated as the average standard deviation of the projections of trials within each class. Finally, the population d’ measure was calculated as *d*^′^ = Δ*μ*/*σ*. To calculate single-neuron measures of discriminability, we calculated for each neuron, the separation (Δ*μ*) as the absolute difference in the average firing rate for the two classes, the noise (*σ*) as the average standard deviation in firing rate for each class, and the discriminability (d’) was calculated as *d*^′^ = Δ*μ*/*σ*. We repeated this calculation for all pairs of food items, for both monkeys, and for 5,000 iterations of sampling trials. The mean and standard deviation of these values across iterations are depicted in Figure 6.

#### Neural discriminability in the natural task

To estimate the neural discriminability in the natural task and its variability, we performed a bootstrap analysis. In each iteration, we selected trials from the natural task with replacement, so that the total number matched the number of trials in the natural task. We then trained a population decoder on these resampled trials as before, and obtained the decoder accuracy using leave-one-out cross-validation. We then calculated the population discriminability, mean separation and noise standard deviation as well as their single neuron counterparts as before. This process was repeated for 5,000 times.

#### Statistical comparisons of neural discriminability

Since the above bootstrap sampling can be performed any arbitrary number of times, standard statistical tests cannot be used to assess statistical significance. Instead, we deemed a comparison (A > B) to be statistically significant if the number of violations of that trend (i.e. A < B) occurred only in a small fraction of the iterations (*α* < 0.05). This statistical significance is reported in Figure 4D-E.

### Event dissimilarity during the natural task (Figure 7)

For each event in the natural task, we calculated the firing rate of each neuron in 0.1 s bins spanning a 0.5 s window around the event onset, and averaged it across all occurrences of the event to create a population activity vector with 128*5 dimensions. To calculate the event dissimilarity between a pair of events, we calculated the correlation distance (1 – *r*) between the population activity vectors of the two events, where *r* is the Pearson’s correlation (Figure 7).

In addition, rather than depicting the event dissimilarity for an event with itself as zero, we calculated its noise level. We took the correlation between the population vectors obtained from the odd and even half of all trials of that event, and corrected it using the Spearman-Brown correction 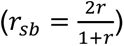. Thus, a large value indicates a noisy event representation, whereas a small value indicates a highly reliable event representation.

### Visualization of event representations in the natural task

To visualize event representations related to food or experimenter events, we performed a multidimensional scaling (*mdscale* function, MATLAB 2022). This procedure identifies 2D coordinates for each event such that the distances in 2D match have the best possible match with the observed event dissimilarities. In the resulting plots, nearby events indicate that they evoked similar neural responses.

## ACKNOWLEDGEMENTS

We thank Marius Peelen & Ashesh Dhawale for valuable discussions, Stanislas Dehaene and Rufin Vogels for valuable comments on the manuscript, Dr. Tim Blanche of White-Matter LLC for technical support with wireless recordings, Jason Joby for helping with code review, Dr. Sebastian Chandu for assistance with implant surgeries, Dr. Sumedh Shastry for help with physiology during surgery and post-surgical care, Dr. Sumedh Shastry & Mr. V. Ramesh for primate facility management and animal care, and Mr. Ravi & Mr. Ashok for animal care.

## AUTHOR CONTRIBUTIONS

Sini Simon: Recording setup, Analysis, Data visualization, Manuscript writing & editing. Specific contributions: Developed hardware & software for wireless brain recordings, tested electrodes, trained monkeys, planned and conducted implant surgeries, participated in troubleshooting recordings, performed data analysis, data visualization, created videos, wrote manuscript and integrated comments from all authors.

Surbhi Munda: Recording setup, Experiment design, Data collection & curation, Analysis, Manuscript writing & editing. Specific contributions: Developed hardware & software for wireless brain recordings, trained monkeys, planned and conducted implant surgeries, participated in troubleshooting recordings, designed experiments, collected data, curated & annotated data, helped with data visualization, performed data analysis, wrote sections of the manuscript, created videos, provided feedback on the manuscript drafts.

Shubhankar Saha: Recording setup, Data Analysis, Manuscript editing. Specific contributions: Developed hardware & software for wireless brain recordings, trained monkeys, planned and conducted implant surgeries, performed data analysis, wrote sections of the manuscript, provided feedback on the manuscript drafts.

Thomas Cherian: Recording setup, Experiment Design. Specific contributions: Developed hardware & template software codes for recording, tested electrodes and implant housing, planned and conducted implant surgeries, participated in troubleshooting recordings, and designed motor tasks.

Georgin Jacob: Recording setup, Manuscript editing. Specific contributions: Developed hardware & template software codes for recording, developed software for automated spike detection and sorting, tested electrodes and implant housing, planned and conducted implant surgeries, participated in troubleshooting recordings, and provided feedback on manuscript drafts.

Jhilik Das: Recording setup, Data Analysis, Manuscript Editing. Specific contributions: Developed hardware & software codes for data curation and analysis, tested electrodes and implant housing, snout trained monkeys, planned and conducted implant surgeries, participated in troubleshooting recording setup, performed data analysis, and provided feedback on manuscript drafts.

Debdyuti Bhadra: Recording setup. Experiment design, Data collection, Data curation, Data visualization and Manuscript editing. Specific contributions: Designed experiments, collected data, helped with hardware and software updates for wireless brain recordings, planned and conducted implant surgeries, provided feedback on manuscript drafts, created videos.

## DATA AVAILABILITY

All data and code required to reproduce the results are publicly available at https://osf.io/3bpyv

## SUPPLEMENTARY MATERIAL

**Figure S1.**
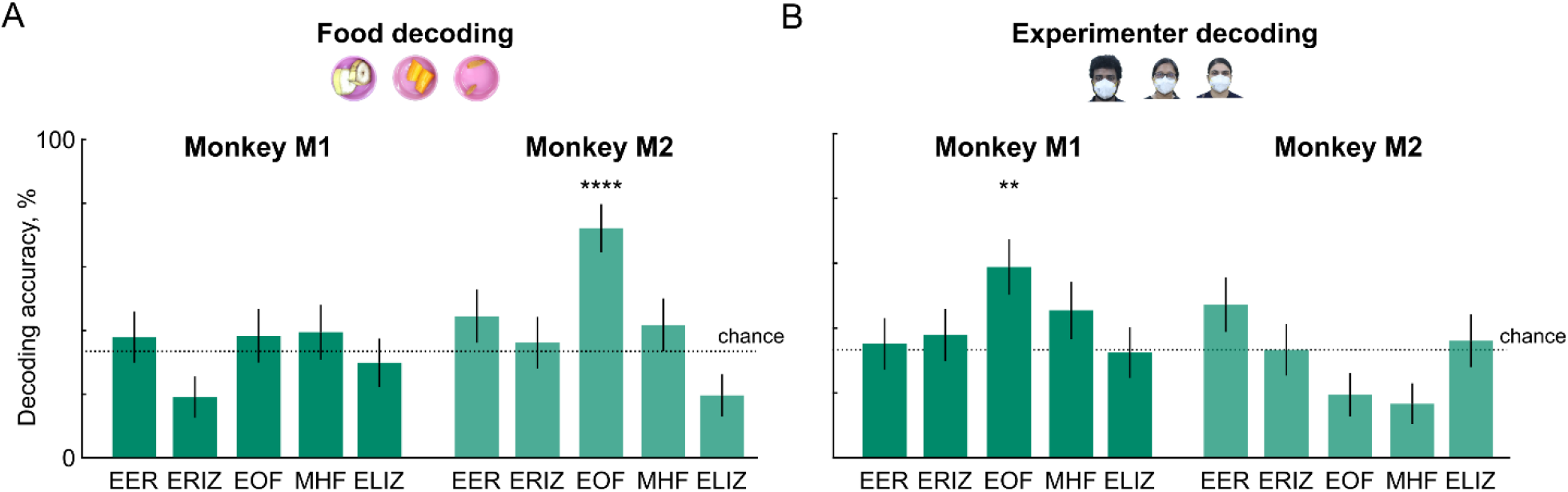
Differences in food & experimenter decoding between monkeys. During the natural task, we noticed subtle differences in the behavior of the two monkeys. While taking the food from the experimenters’ hand, monkey M1 looked at the experimenter, while monkey M2 looked at the food. We therefore wondered whether this would be reflected in the decoding of food and experimenter identity in the two monkeys. To investigate this issue, we trained food and face decoders in the fixation task and tested them around key events in the natural task. Event abbreviations: EER: Experimenter Entered Room; EOF: Experimenter Offered Food; MHF: Monkey Held Food. (A) Food decoding accuracy (mean ± s.e.m across events) for key behavioral events in the natural task for Monkey M1 (*left*) and M2 (*right*). Asterisks represent statistical significance (*** is p < 0.0005, chi-squared test comparing observed counts with those predicted by chance). Consistent with our observations, we found food decoding to be best in monkey M1 when it was holding food (MHF) and in monkey M2 when experimenter offered food (EOF). Event abbreviations: EER: Experimenter Entered Room; EOF: Experimenter Offered Food; MHF: Monkey Held Food. (B) Experimenter decoding accuracy (mean ± s.e.m across events) for key behavioral events in the natural task for monkey M1 (*left*) and M2 (*right*). Consistent with our observations, we found experimenter decoding to be best in monkey M1 when experimenter offered food (EOF) and in monkey M2 when the experimenter entered room (EER). *[Face images shown are those of the authors: D.B., S.M. and S.S.]*

**Figure S2.**
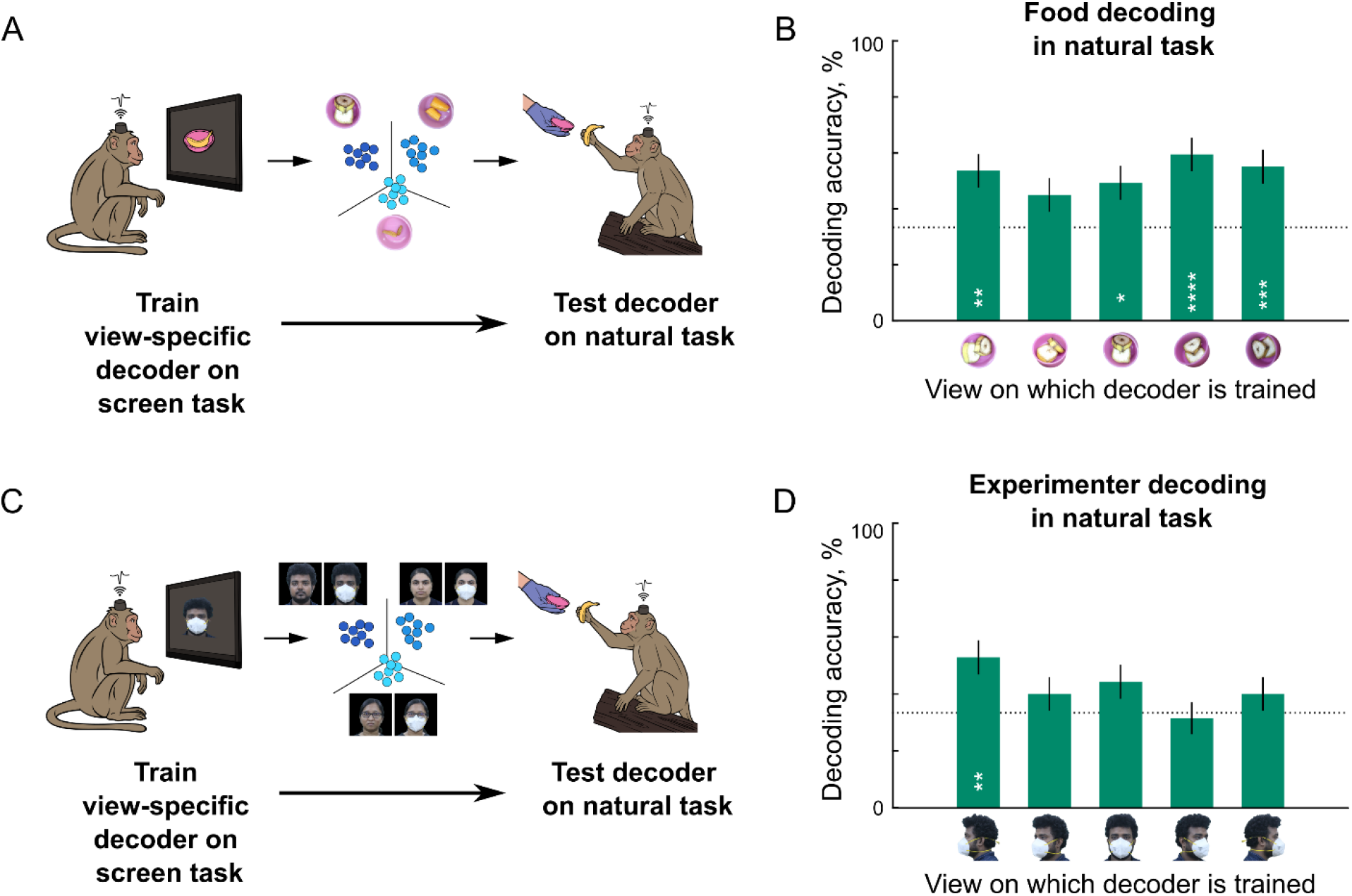
View-specific food & experimenter decoding in the natural task. (A) Schematic of a linear decoder trained on each view of food box images in the fixation task and then tested in the natural task at the time of Monkey Held Food (MHF) for M1, and Experimenter Offered Food (EOF) for M2, when the monkeys viewed the food (based on our own observations during recording). (B) Decoding accuracy for each view-specific food decoder. It can be seen that food decoding was more accurate at some views compared to others. These are close to the food box views seen by the monkeys during the natural task. (C) Same as (A) but for experimenter decoding. Classifiers were trained on specific views of experimenter faces (both with and without face mask) and tested in the natural task at the time of Experimenter Offered Food (EOF) for M1, and Experimenter Entered Room (EER) for M2, when the monkeys looked at the experimenter (based on our own observations during recording). *[Face images shown are those of the authors: D.B., S.M. and S.S]* (D) Same as (B) but for experimenter decoding. It can be seen that face decoding accuracy was most accurate for side views of the experimenters’ face. These are close to the face view seen by the monkeys in the natural task. *[Face images shown are those of one of the authors, D.B.]*

**Figure S3.**
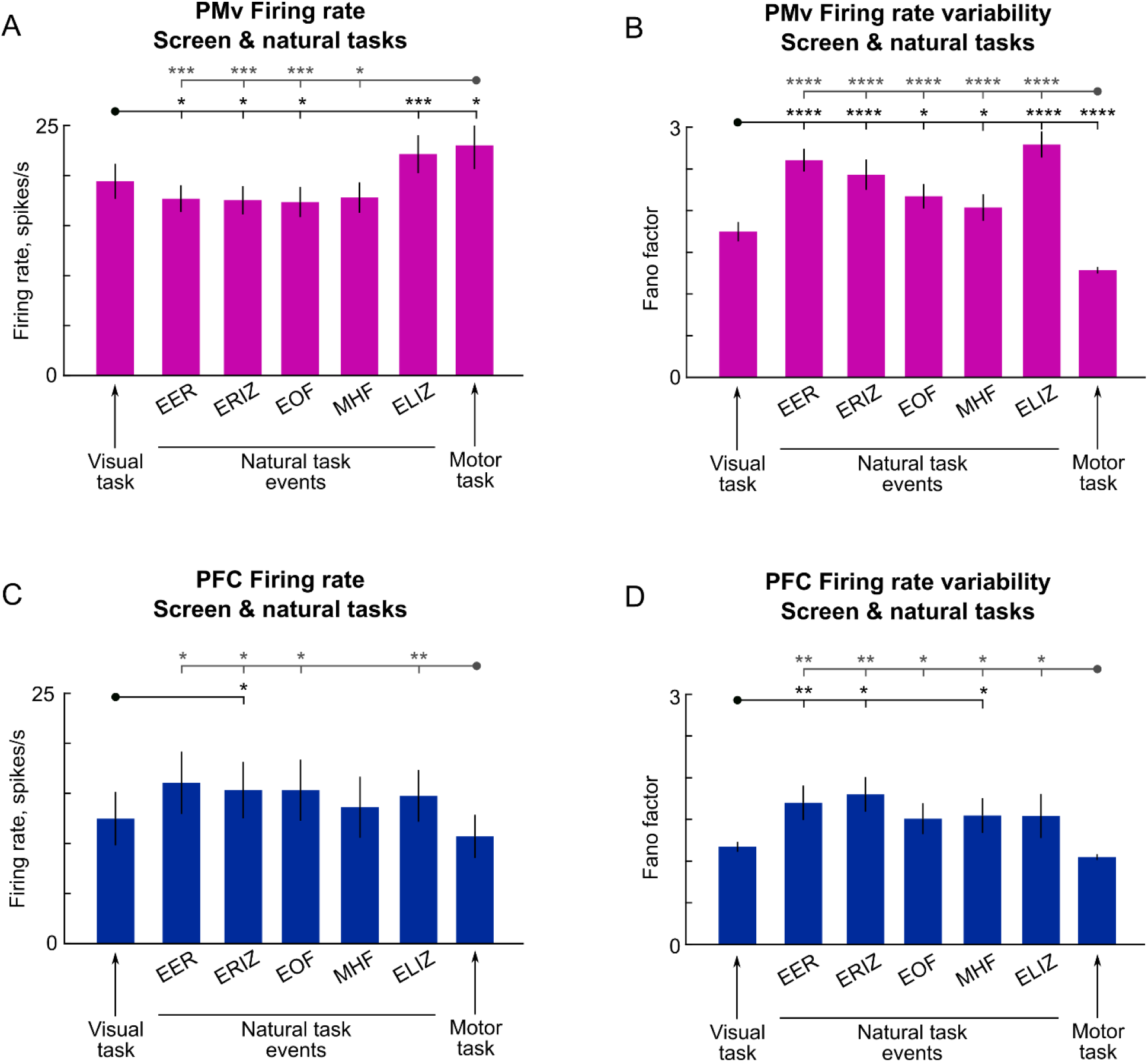
Variability of PMv and PFC during natural and screen tasks. (A) Average firing rate across PMv neurons (pooled across monkeys) in the screen & natural tasks in equivalent time windows (same as Figure 4A). Asterisks indicate statistical significance of the corresponding comparison (* is p < 0.05, ** is p < 0.005, etc; sign-rank test across firing rates of 93 motor neurons; comparisons shown only between each screen task with all events in the natural task). It can be seen that neural responses were highest for the motor task compared to most natural task events. (B) Variability of PMv neurons in screen and natural tasks, calculated using the average Fano factor. It can be seen that neural variability was generally higher for the natural task events compared to the screen tasks. (C) Same as (A) but for PFC neurons in monkey M1 (n =16). PFC firing rates are larger in the natural task compared to the screen tasks. (D) Same as (B) but for PFC neurons in monkey M1. PFC shows more variability for many events in the natural task compared to the screen tasks.

**Figure S4.**
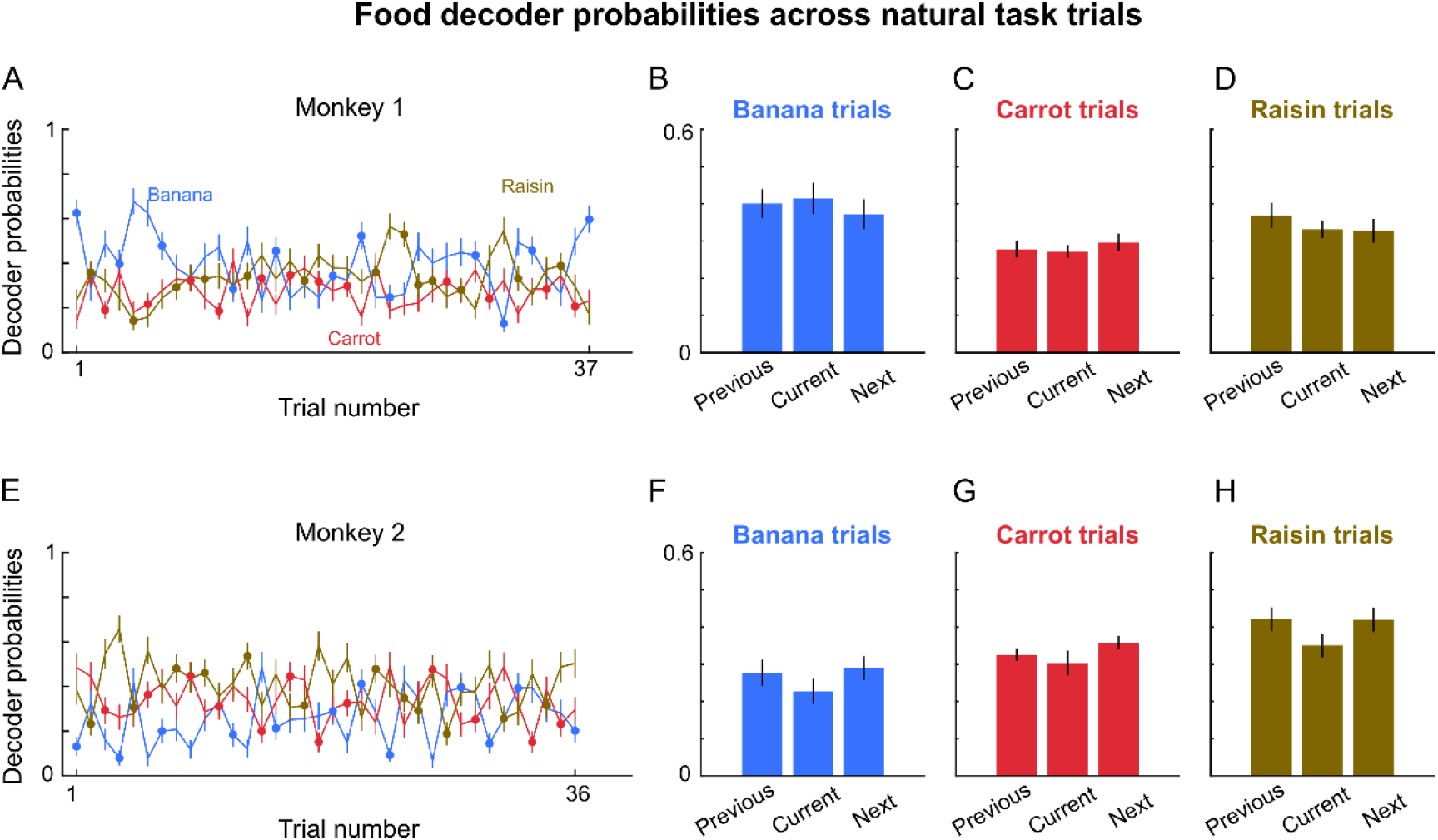
Food decoding dynamics across trials in the natural task. Since the monkeys received food in a particular sequence across trials, we wondered whether they started to anticipate specific food items as the trials progressed. Trials in the natural task were designed such that no two consecutive trials had the same food or experimenter. To explore this issue, we calculated decoder probability of each food for each trial at the time of experimenter entering the room, for the decoder trained on the fixation task trials and tested on the natural task trials. (A) Decoder probability in monkey M1 for banana (*blue*), carrot (*red*) and raisin (*brown*) for each trial in the natural task, with dots indicating trials where that particular food was given. (B) Average decoder probability for the trial on which banana was given, the previous trial, and the next trial. It can be seen that banana probability was high, reflecting anticipation. Since the trials were designed such that consecutive trials do not involve the same food, the absence of a banana in the previous trials is a predictor of the banana being offered on the current trial. The s.e.m is calculated across repeats of the food condition. (C) Same as (B) but for carrot trials. Here we do not see any anticipatory response. (D) Same as (B) but for raisin trials. Here we see a higher probability for raisin in the trial before the raisin was offered. This too is possible, since the raisin trial was preceded by a non-raisin trial and could have led to an expectation on part of the monkey. (E) Same as (A) but for monkey M2. (F) Same as (B) but for monkey M2. Banana decoding probability was higher for the trial immediately preceding the banana, presumably because the preceding trials did not involve a banana. It was higher for the trial immediately after, presumably because monkey M2 expected a repeat trial. (G) Same as (F) but for carrot trials, with the same decoding patterns as banana trials. (H) Same as (F) but for raisin trials. with the same decoding patterns as banana trials.

**Figure S5.**
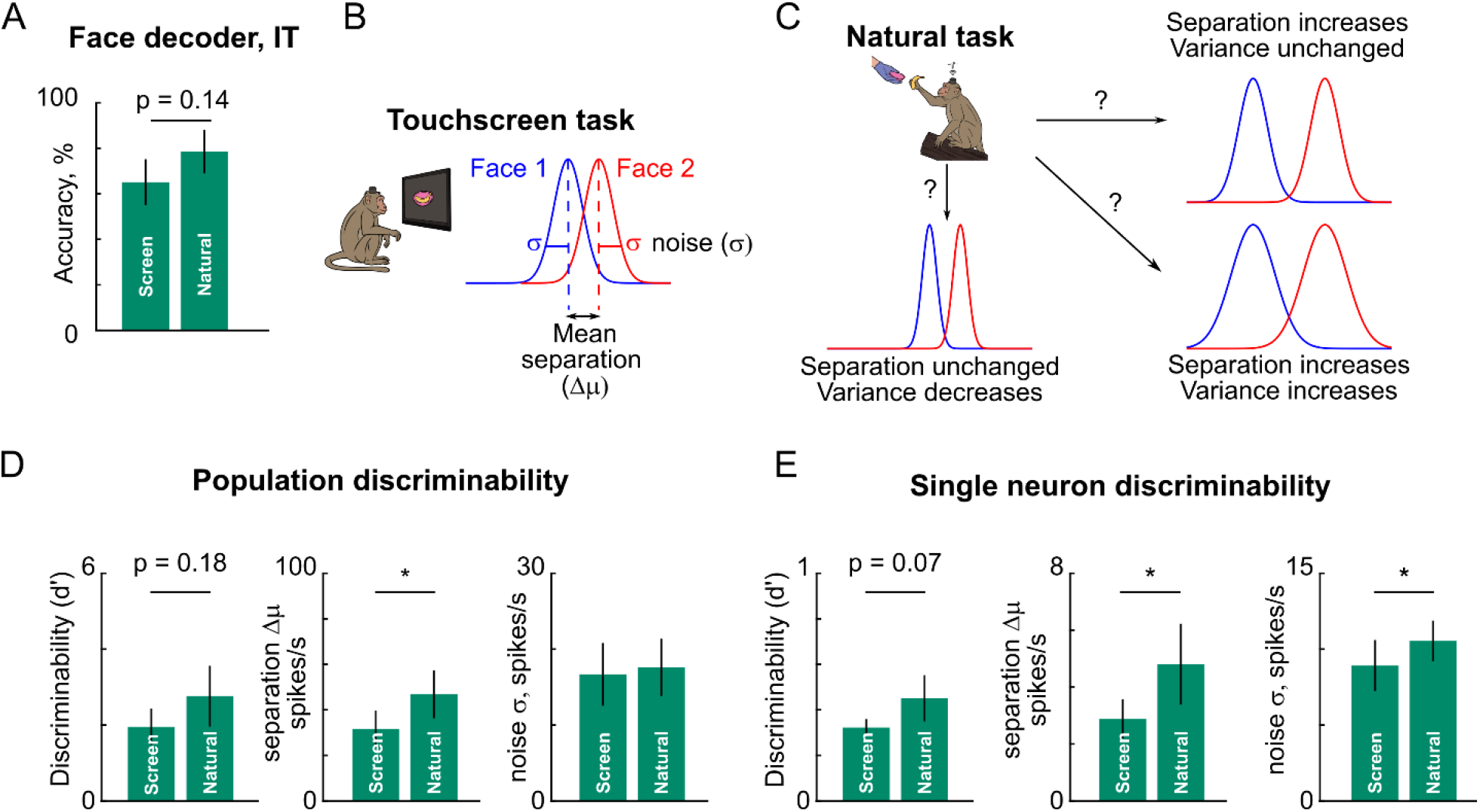
Fidelity of face information in screen & natural tasks. (A) Face decoder accuracy for screen and natural tasks for IT neurons (mean ± sd across bootstrap samples). Decoder accuracy was calculated by bootstrap sampling of equal numbers of trials in the screen and natural tasks (see Methods). The decoder was trained in window of 0 to 0.4 s from image onset in fixation and from event of interest in natural task. Asterisks represent statistical significance, calculated as the frequency of violations of the trend across bootstrap samples (* is p < 0.05, ** is p < 0.005). (B) Schematic showing the distribution of projections of population activity along the linear classifier vector for Face 1 (*blue*) vs Face 2 (*red*). To evaluate the fidelity of information in each task, we compared the mean separation between the classes (Δ*μ*), the noise within each class (*σ*) and the overall discriminability (*d*^′^ = Δ*μ*/*σ*). (C) Schematic showing three possible explanations for increased decoding accuracy in the natural task compared to the fixation task: either the separation increases without change in noise, separation is unchanged with a decrease in noise, or both separation and noise increase. (D) Population discriminability, separation and noise levels for IT neurons, with error bars representing standard deviation across bootstrap samples. For details of each calculation, see Methods. Asterisks represent statistical significance, calculated as the frequency of violations in the depicted trend across bootstrap samples (* is p < 0.05). It can be see that the discriminability is slightly larger in the natural task, driven by an increased separation without an appreciable increase in noise. (E) Same as (F) but using single-neuron analyses. For each neuron we calculated the discriminability, separation and noise standard deviation (see Methods). Asterisks represent statistical significance as before. It can be seen that single neuron discriminability is larger in the natural task, and this is driven by an increase in separation that is larger than the increase in noise levels.

**Figure S6.**
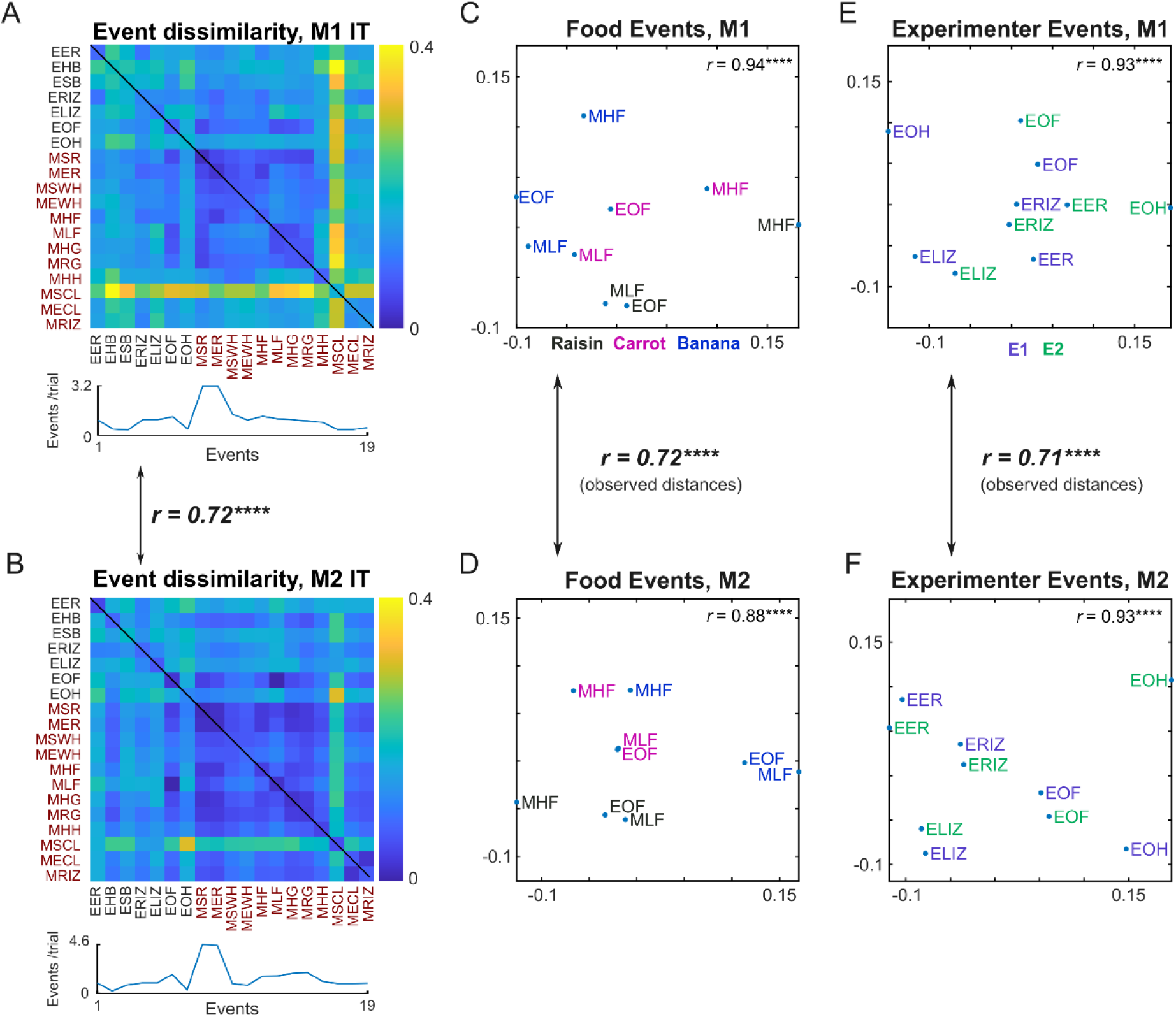
Similar event representations in M1 & M2 IT neurons. (A) *Top:* Neural dissimilarity in monkey M1 IT neurons between all pairs of annotated events in the natural task. *Bottom:* Average number of occurrences per trial of each event. *Event abbreviations:* EER, Experimenter Entered Room; ESB, Experimenter Show Box; EHB, Experimenter Hide Box; ERIZ, Experimenter Reached Interaction Zone, ELIZ, Experimenter Left Interaction Zone; EOF, Experimenter Offered Food; EOH, Experimenter Offered Hand; MER-Monkey End Reach; MSR, Monkey Start Reach; MSWH, Monkey Start Withdraw Hand, MEWH, Monkey Stop Withdraw Hand; MHF, Monkey Held Food, MLF, Monkey Look At Food; MHG, Monkey Held Grill; MRG, Monkey Released Grill; MHH, Monkey Held Hand; MSCL, Monkey Start Climb Log; MECL, Monkey End Climb Log; MRIZ, Monkey Reached Interaction Zone. (B) Same as (A) but for Monkey M2 IT neurons. The correlation between the event dissimilarities in M1 & M2 is indicated (**** is p < 0.00005). Thus, key events in the natural task have a similar representation in IT neurons of both monkeys. (C) Multidimensional scaling plot of event dissimilarities in monkey M1 IT for a subset of events that involved food, but colored separately for by food (*blue*, banana; *magenta*, carrot; *black*, raisin). (D) Same as (E) but for monkey M2. The correlation between the event dissimilarities in M1 & M2 is indicated, spanning panel C (**** is p < 0.00005). Thus, key food-related events in the natural task have a similar representation in IT neurons of both monkeys. (E) Multidimensional scaling plot of event dissimilarities in monkey M1 IT for a subset of events that involved the human experimenter, but colored separately by experimenter (*blue*, E1; *green*, E2). In the plot, events are nearby if they elicited similar neural activity. The correlation at the top shows the match between pairwise event dissimilarities and the 2D distances in the plot (asterisks represent statistical significance: **** is p < 0.00005). (F) Same as (C) but for monkey M2. The correlation between the event dissimilarities in M1 and M2 is indicated spanning panel E (**** is p < 0.00005). Thus, key experimenter-related events in the natural task have a similar representation in IT neurons of both monkeys.

**Figure S7.**
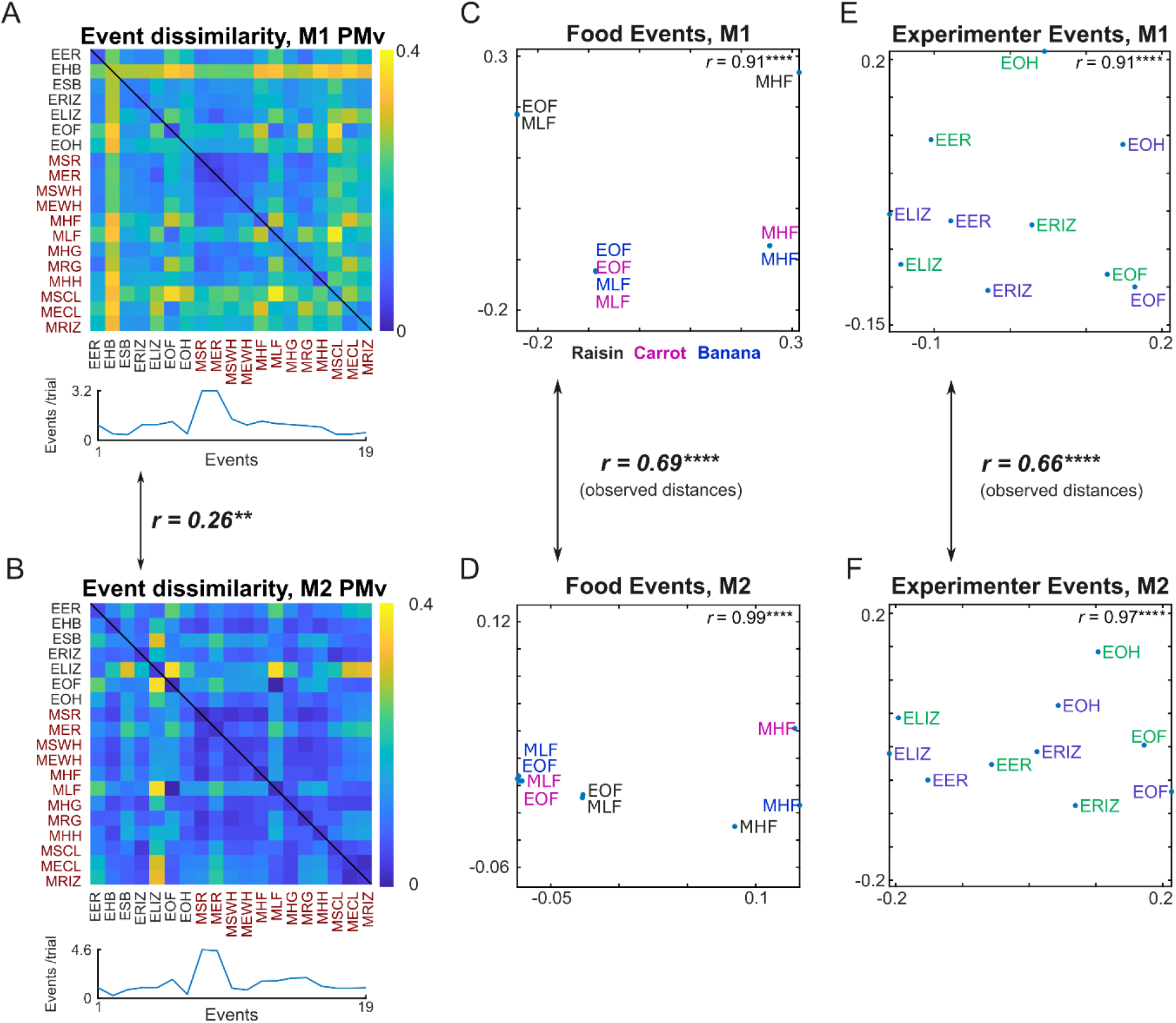
**Similar event representations in M1 & M2 PMv neurons. All conventions are as in Figure S6.**

